# A Broadly Conserved Deoxycytidine Deaminase Protects Bacteria from Phage Infection

**DOI:** 10.1101/2021.03.31.437871

**Authors:** Geoffrey B. Severin, Brian Y. Hsueh, Clinton A. Elg, John A. Dover, Christopher R. Rhoades, Alex J. Wessel, Benjamin J. Ridenhour, Eva M. Top, Janani Ravi, Kristin N. Parent, Christopher M. Waters

**Affiliations:** Department of Biochemistry and Molecular Biology, Michigan State University, East Lansing, Michigan, USA, 48824; Department of Microbiology and Molecular Genetics, Michigan State University, East Lansing, Michigan, USA, 48824; Department of Biological Sciences, Institute for Bioinformatics and Evolutionary Studies, Bioinformatics and Computational Biology Program, University of Idaho, Moscow, Idaho, USA, 83844; Department of Mathematics and Statistical Sciences, University of Idaho, Moscow, Idaho, USA, 83844; Department of Pathobiology and Diagnostic Investigation, Michigan State University, East Lansing, Michigan, USA, 48824

**Keywords:** cytidine deaminase, phage, genomic island, toxin-antitoxin, thymineless death, APOBEC

## Abstract

The El Tor biotype of *Vibrio cholerae* is responsible for perpetuating the longest cholera pandemic in recorded history (1961-current). The genomic islands VSP-1 and -2 are two understudied genetic features that distinguish El Tor from previous pandemics. To understand their utility, we calculated the co-occurrence of VSP genes across bacterial genomes. This analysis predicted the previously uncharacterized *vc0175*, herein renamed **d**eoxycytidylate **d**eaminase ***V****ibrio* (*dcdV*), is in a gene network with *dncV*, a cyclic GMP-AMP synthase involved in phage defense. DcdV consists of two domains, a P-loop kinase and a deoxycytidylate deaminase, that are required for the deamination of dCTP and dCMP, inhibiting phage predation by corrupting cellular nucleotide concentrations. Additionally, DcdV is post-translationally inhibited by a unique noncoding RNA encoded 5’ of the *dcdV* locus. DcdV homologs are conserved in bacteria and eukaryotes and our results identify *V. cholerae* DcdV as the founding member of a previously undescribed bacterial phage defense system.

## INTRODUCTION

*Vibrio cholerae*, the etiological agent responsible for the diarrheal disease cholera, is a monotrichous, Gram-negative bacterium found ubiquitously in marine environments [1]. There have been seven recorded pandemics of cholera, beginning in 1817, and the fifth and sixth pandemics were caused by strains of the classical biotype. The seventh pandemic, which began in 1961 and continues today, was initiated and perpetuated by circulating strains of the El Tor biotype. Numerous phenotypic and genetic characteristics are used to distinguish the classical and El Tor biotypes [2]. It is hypothesized that El Tor’s acquisition of two unique genomic islands of unknown origins, named the Vibrio Seventh Pandemic Islands 1 and 2 (VSP-1 and 2) [3], played a pivotal role in El Tor’s evolution to pandemicity and the displacement of the classic biotype in modern cholera disease [4].

Combined, VSP-1 and VSP-2 encode ∼36 putative open reading frames (ORFs) within ∼39 kb (Figs. 1A and S1B) [3, 5–7]. While the majority of the genes in these two islands remain to be studied, it is hypothesized that the biological functions they encode may contribute to environmental persistence [8] and/or the pathogenicity [9] of the El Tor biotype. In support of this idea, VSP-1 encodes a phage defense system encompassing the genes *dncV, capV*, *vc0180* and *vc0181* called the cyclic-oligonucleotide-based antiphage signaling system (CBASS) [10] (Fig. 1A). CBASS limits phage invasion of bacterial populations via a process termed abortive replication whereby upon phage infection cyclic GMP-AMP (cGAMP) synthesis by DncV activates cell lysis by stimulating the phospholipase activity of CapV [10, 11]. During our search for VSP-1 and 2 gene networks, we determined that the gene *vc0175,* renamed herein as **d**eoxy**c**ytidylate **d**eaminase in ***V****ibrio* (*dcdV*), cooccurs in bacterial genomes with *dncV*, suggesting a common function.

**Fig. 1:**
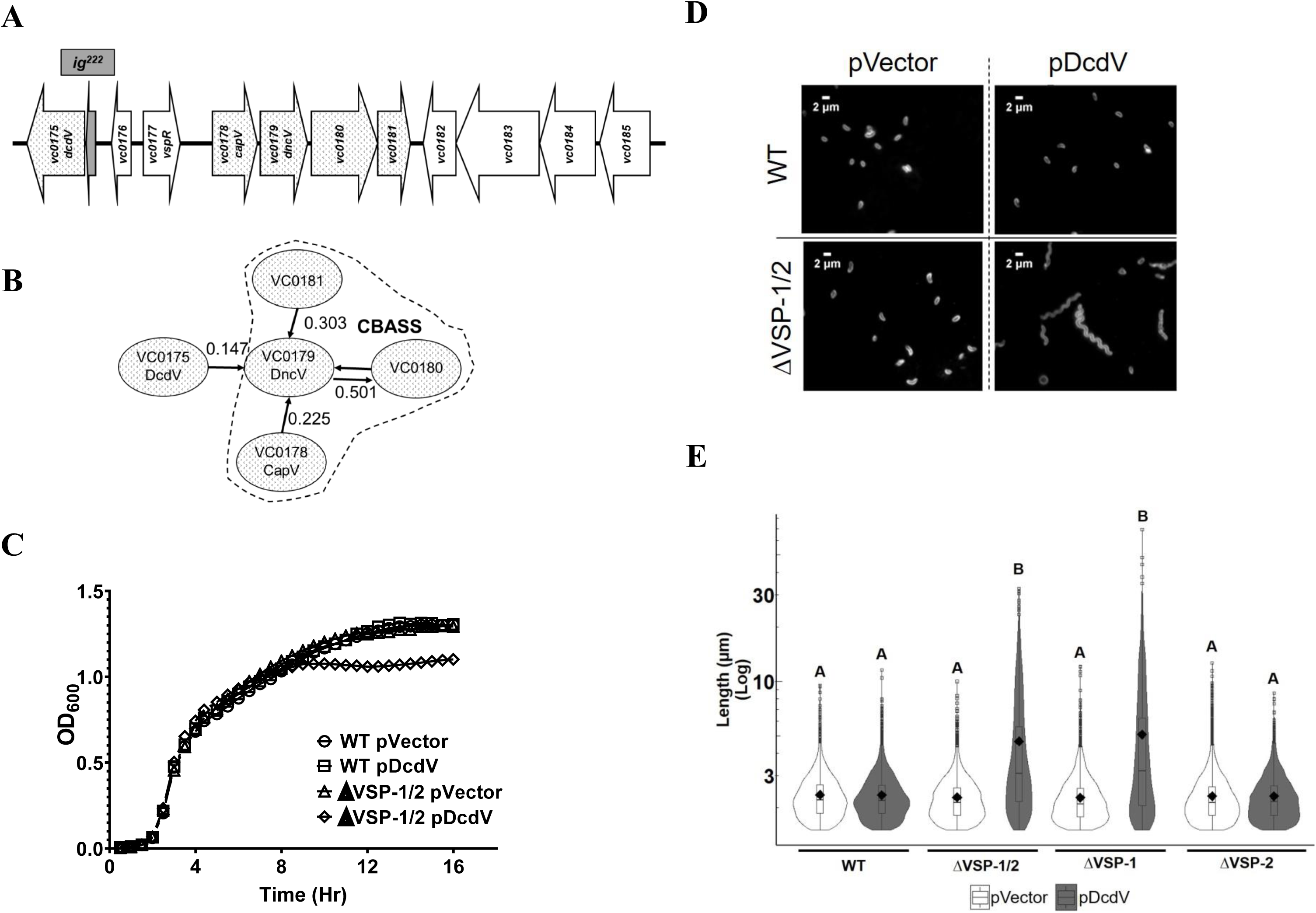
DcdV promotes filamentation in *V. cholerae* in the absence of VSP-1. (**A**) Cartoon schematic of VSP-1 and (**B**) the Correlogy gene network prediction for *dncV* where arrows show the highest partial correlation *W_ij_* each individual VSP-1 gene has to another. (**C**) Growth of WT *V. cholerae* and ΔVSP-1/2 strains with the vector or pDcdV. Data represent the mean ± SEM, *n*=3. (**D**) Representative images of WT and ΔVSP-1/2 strains with the vector or pDcdV. (**E**) Violin plots of cell length distributions of WT, ΔVSP-1/2, ΔVSP-1, and ΔVSP-2 strains with the vector or pDcdV: summary statistic for this and all following violin plots are mean (diamonds), median (horizontal black line), interquartile range (box), and data below and above the interquartile range (vertical lines). Different letters indicate significant differences (*n*=3) at p < 0.05, according to Tukey’s post-hoc test.

We show that *dcdV,* exhibits deoxycytidylate deaminase (DCD) activity, catalyzing the deamination of free deoxycytidine monophosphate (dCMP) substrates to form deoxyuridine monophosphate (dUMP) and is part of the broader Zn-dependent cytosine deaminase (CDA) family of enzymes [12–14]. The activity of DCD enzymes play a vital role in the de novo synthesis of deoxythymidine triphosphate (dTTP) by supplying the dUMP required by thymidylate synthase (TS) to form deoxythymidine monophosphate (dTMP) [12]. CDA enzymes belonging to the APOBEC (Apolipoprotein B mRNA editing enzyme catalytic polypeptide-like) family also play an important role in viral immunity in higher organisms where their catalytic activity is utilized for the deamination of nucleic acids rather than free nucleotide substrates to restrict several types of viruses, such as retroviruses, and retroelements [15–19].

A primary challenge faced by lytic phage is to rapidly replicate many copies of its genome, which requires sufficient nucleotide substrates [20]. During DNA phage infection, total DNA within a bacterium can increase 5-10 fold, illustrating the vast amount of DNA replication that must occur in a short window of time [21, 22]. To accomplish this feat, invading DNA phage often corrupt the delicate balance of enzymatic activity across a host’s deoxynucleotide biosynthetic pathways by deploying their own DCD, dUTPase, TS, and ribonucleotide reductase to ensure the appropriate ratio and abundance of deoxyribonucleotides [23–27].

Here we show that DcdV is a dual domain protein consisting of a putative N-terminal P-loop kinase (PLK) and C-terminal DCD domain, and this novel domain architecture is present across the tree of life. Overexpression of DcdV promotes cell filamentation, which has hallmarks of nucleotide starvation resembling thymine-less death (TLD) toxicity [28–31]. Our results demonstrate that ectopic expression of DcdV indeed corrupts the intracellular concentrations of deoxynucleotides and this activity protects bacteria from phage infection. Moreover, we demonstrate that DcdV activity is negatively regulated by a non-coding RNA encoded 5’ of the *dcdV* locus [renamed herein as **D**cdV **i**nsensitivity **f**actor in ***V****ibrio* (*difV*)]. Furthermore, *dcdV-difV* systems are widely encoded in bacteria and we show that a subset of them function similarly, establishing cytidine deaminase enzymes as antiphage defense systems in bacteria.

## RESULTS

### *dncV* and *dcdV* co-occur in bacterial genomes

To help identify functional interactions within the largely unclassified VSP-1 & 2 genes, VSP island genes were classified into putative “gene networks” or sets of genes that form a functional pathway to accomplish a biological task. Since gene networks often share deep evolutionary history among diverse taxa, we hypothesized that the set of genes in a gene network would co-occur together in the genomes of diverse taxa at a higher frequency than chance alone would predict. Our software package was named ‘Correlogy’ inspired by [32] and is described in detail in the materials and methods.

We calculated a Pearson correlation followed by a partial correlation correction between each of the VSP island genes from the same island across the sequenced bacterial domain.

This resulting partial correlation correction ”*w_ij_*” has an output normalized to a range of -1 to 1, with a *w_ij_* of -1 revealing homologs of genes *i* and *j* never occur in the same species as opposed to a value of 1 in which homologues of genes *i* and *j* always co-occur in the same species. Previous research using well-classified *Escherichia coli* gene networks showed that partial correlation values *w_ij_* > 0.045 were highly correlated with shared biological functions [32]. Using the above-mentioned approach, we calculated a partial correlation value *w_ij_* for all genes *i* to *j* in VSP-1 (Supplemental File 1) and VSP-2 (Supplemental File 2). We generated a visualization of the Maximum Relatedness Subnetworks (MRS) showing the single highest *w_ij_* value for each VSP gene (Figs. 1B, S1A, 1B).

One of our VSP-1 gene networks centered on *dncV* and identifies the experimentally validated CBASS anti-phage system (Fig. 1B) [10]. Curiously, the putative deoxycytidylate deaminase encoded by *vc0175,*which we renamed *dcdV*, was also found to co-occur with *dncV* (*w_ij_* = 0.147) but not with any of the other CBASS members (*w_ij_* < 0.045) (Fig. 1B). Recognizing that co-occurrence of *dncV* with *dcdV* may indicate a shared or common biological function, we sought to understand the biological activity of *dcdV*.

### Ectopic expression of *dcdV* induces cell filamentation in the absence of VSP-1

To assess the function of DcdV, we generated growth curves in both wild type (WT) *V. cholerae* and a double VSP island deletion strain (ΔVSP-1/2) over-expressing *dcdV* (pDcdV) or vector control (pVector). DcdV overexpression did not impact WT growth but did reduce growth yield in the ΔVSP-1/2 background (Fig. 1C). We evaluated the cellular morphology of WT and ΔVSP-1/2 strains after overexpression of DcdV and observed expression from pDcdV in the ΔVSP-1/2 background yielded filamentous cell morphologies, suggesting these cells have a defect in cell division that manifests in a reduced growth yield (Fig. 1D). We performed the same image analysis in single island mutants (ΔVSP-1 and ΔVSP-2) and found that the mean cell length increased significantly upon DcdV overexpression only in cells lacking VSP-1 (Fig. 1E).

Likewise, overexpression of pDcdV in a laboratory strain of *E. coli* also induced cell filamentation that was inhibited by provision of a single copy cosmid containing VSP-1 (pCCD7) but not the vector cosmid control (pLAFR) (Figs. S2A and S2B). The spiral nature of *V. cholerae* filaments (Fig. 1D) is due to the natural curvature of *V. cholerae* mediated by *crvA* [33, 34]. Taken together, these results indicated that DcdV overexpression severely impacts cell physiology in the absence of VSP-1.

### DifV is encoded immediately 5’ of the *dcdV* locus in VSP-1

To identify the negative regulator of DcdV activity encoded in VSP-1, we generated partial VSP-1 island deletions and quantified cell filamentation following DcdV expression. Three sections of VSP-1; *dcdV-vc0176*, *vspR-vc0181,* and *vc0182-vc0185,* were individually deleted. Of the three partial VSP-1 deletion strains, expression of pDcdV only induced filamentation in the Δ*dcdV*-*vc0176* mutant (Fig. 2A). Individual gene deletion mutants of *dcdV* and *vc0176* maintained WT cell morphology following expression of DcdV (Fig. 2B), suggesting the 504 nt intergenic region between *dcdV* and *vc0176* is the source of DcdV inhibition. We identified a 222 nucleotide (nt) open reading frame we named *ig^222^* encoded in the same orientation immediately 5’ of *dcdV* as a possible candidate for the DcdV regulation (Fig. 1A).

**Fig. 2:**
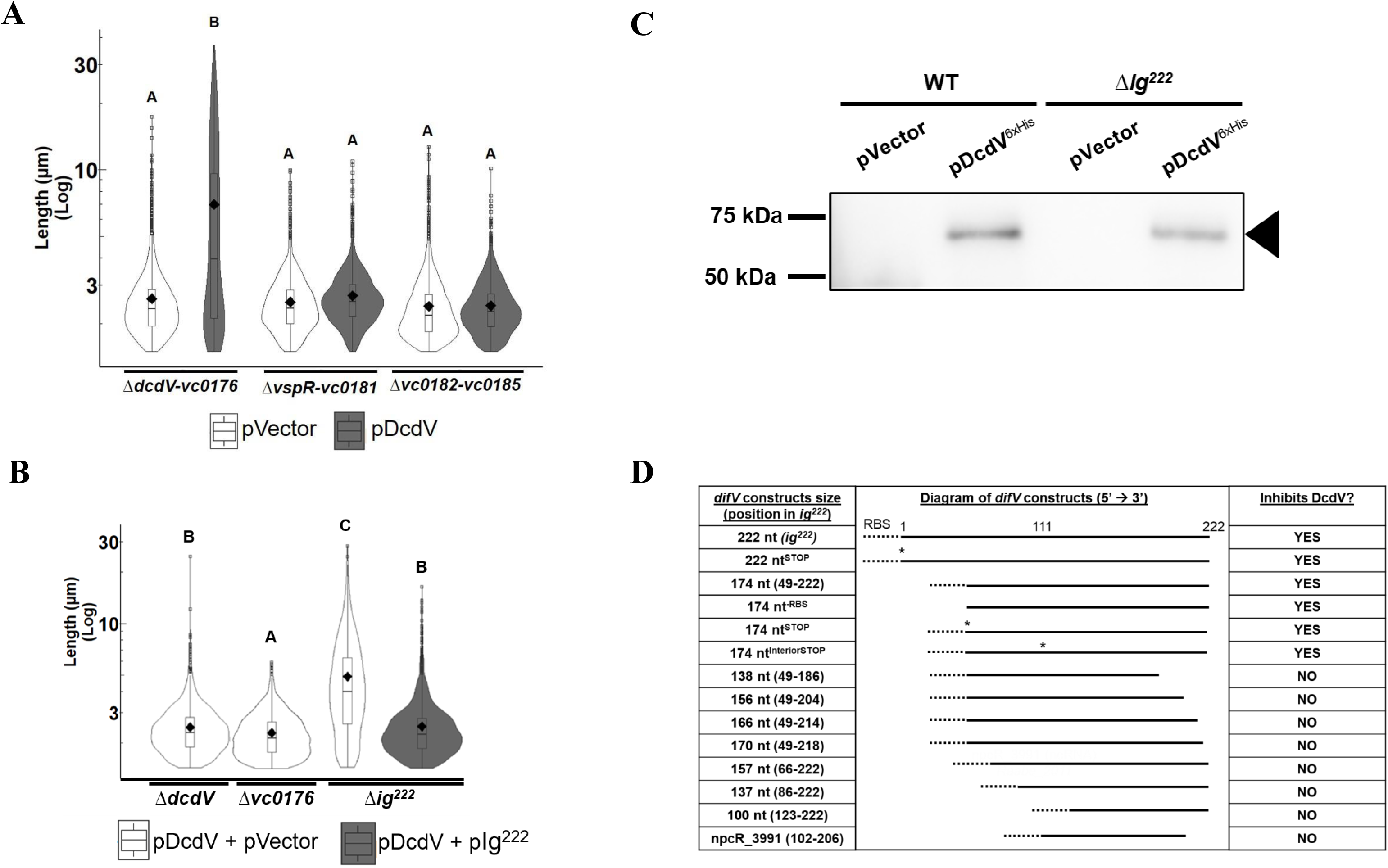
DifV is a sRNA that post-translationally regulates DcdV. (**A**) Distribution of cell lengths measured from three biological replicates of gene deletions within VSP-1 or (**B**) individual gene deletions as indicated containing vector or pDcdV grown in the presence of 100 µM IPTG for 8 h. Different letters indicate significant differences (*n*=3) at *p* < 0.05, according to Tukey’s post-hoc test. (**C**) Representative anti-6x His antibody Western blot of whole cell lysates from *V. cholerae* WT and Δ*ig^222^* cultures maintaining vector or pDcdV^6xHis^. Analysis was performed in triplicate biological samples. Black triangle corresponds to DcdV^6xHis^ (60.6 kDa). (**D**) Table of various *difV* constructs expressed in Δ*ig^222^* under a P_tac_-inducible promoter with a non-native ribosomal binding site (RBS, denoted by dotted line). DcdV induced filamentation in the presence of these *difV* constructs was assessed using fluorescence microscopy in biological triplicate cultures. “**\***” indicates a stop codon introduced in place of a putative start codon.

Overexpression of DcdV in the Δ*ig^222^* mutant led to cell filamentation (Fig. 2B). Furthermore, complementation of *ig^222^* co-expressed from a second plasmid in the *Δig^222^* strain prevented DcdV induced filamentation (Fig. 2B). We conclude that *ig^222^* contains the necessary genetic components for inhibiting DcdV activity and refer to this negative regulator as DifV for **D**cdV **i**nsensitivity **f**actor in ***V****ibrios*.

As *dcdV* and *dncV* cooccur in a gene network (Fig. 1B), we hypothesized that the role of DncV was to inactivate DifV, leading to the liberation of DcdV activity. However, co-expression of both DncV and DcdV did not liberate DcdV activity as these cells were not filamentous (Fig. S3). The Δ*ig^222^* mutant is not filamentous in the absence of pDcdV expression which is likely due to a polar effect originating from the deletion of *ig^222^*. Indeed, *dcdV* expression was reduced at all growth phases in the Δ*ig^222^* mutant (Fig. S4).

### DifV is an sRNA that post-translationally regulates the activity of DcdV

The fact that DifV inhibits DcdV expressed from a plasmid with exogenous transcription and translation start sites suggests DifV regulates DcdV at a post-translational level. To test this hypothesis, we expressed a *dcdV* C-terminal 6x histidine tagged construct (DcdV^6xHIS^) in WT and Δ*ig^222^ V. cholerae* and probed for the cellular abundance of DcdV^6xHIS^ using Western blot (Fig. 2C). When this tagged DcdV is expressed, Δ*ig^222^* manifest a filamentation phenotype while the WT strain does not, indicating the 6x histidine tag does not change the activity of DcdV nor does it inhibit the ability of DifV to regulate DcdV (Fig. S5). Despite the lack of filamentation in the WT strain, the cellular abundance of DcdV^6xHIS^ was slightly greater than Δ*ig^222^* with an average signal intensity ratio WT:Δ*ig^222^*) of 1.5 ± 0.3 across three biological replicates, although this difference was not statistically significant. This result connotes that DifV limits DcdV activity after it has been translated and not by reducing the abundance of DcdV.

Given that DifV regulates the activity of DcdV at the post-translational level, we wondered if DifV was a small peptide or an untranslated small regulatory RNA (sRNA). Mutation of the *ig^222^* rare CTG start codon to a TAG stop codon (222 nt^STOP^) did not abrogate the ability of this construct to inhibit DcdV activity in trans when co-expressed in the Δ*ig^222^* strain (Fig. 2D).

We then examined a 174 nt ORF completely encoded within *ig^222^* (174 nt) and found it was also sufficient to prevent DcdV induced filamentation (Fig. 2D). Additionally, expression of this 174 nt ORF from constructs either lacking a ribosome binding site (174 nt^-RBS^) or where the native ATG start codon was mutated to a TAA stop codon (174 nt^STOP^) each retained the ability to inhibit DcdV activity (Fig. 2D). We also identified an ATG start codon on the interior of the 174 nt ORF corresponding to an alternative reading frame and mutation of this interior start codon to a TAA stop codon (174 nt^InteriorSTOP^) also failed to abrogate DifV inhibition of DcdV activity (Fig. 2D). Together, these results suggest that translation of a gene product originating from within *ig^222^* is not necessary for DifV activity.

To identify the minimum functional size of *difV* we further truncated this 174 nt segment from both the 5’ and 3’ ends and found that removal of either 18 bp from the 5’ end or 4 bp from the 3’ end was sufficient to abolish DifV activity (Fig. 2D). Additionally, expression in trans of npcR_3991 [35], a 104 nt non-coding RNA of unknown function contained within *ig^222^*, was also unable to inhibit DcdV filamentation (Fig. 2D). Collectively, these results suggest that DifV is a regulatory RNA that is between 152 to 174 nt long encoded 5’ of the *dcdV* locus, and we will therefore refer to the 174 nt locus as *difV* for the remainder of these experiments.

### DifV and DcdV constitute a two gene operon that resembles a Toxin-Antitoxin System

The genomic orientation and proximity of *difV* to *dcdV* suggest they may constitute an operon and two previous genome-wide transcriptional start site (TSS) analyses previously identified a common putative TSS 5’ of *difV* [36, 37]. To test if *difV* and *dcdV* are indeed expressed as an operon, we performed diagnostic PCR with primers located within *difV* and *dcdV* on cDNA generated from both WT and Δ*ig^222^* RNA (Fig. 3A). As expected, *dcdV* was detected in the cDNA generated from each strain while *difV* was only amplified using the WT cDNA template (Fig. 3B). The presence of an 839 nt PCR product amplified using primers spanning *difV* to *dcdV* from the WT cDNA template, that was not present with *Δig^222^* cDNA, confirmed that both genes are present on a shared transcript (Fig. 3B). Additionally, we quantified the relative abundance of *difV* and *dcdV* RNA using qRT-PCR and found the *difV* locus was approximately 40-, 20-, and 60-fold more abundant than *dcdV* at early exponential, late exponential, and stationary phases, respectively (Fig. 3C). While having several unique features, the co-transcription of *difV* and *dcdV* and the post-translational regulation of DcdV activity by the abundant sRNA DifV resembles Type III Toxin-Antitoxin systems [38].

**Fig. 3:**
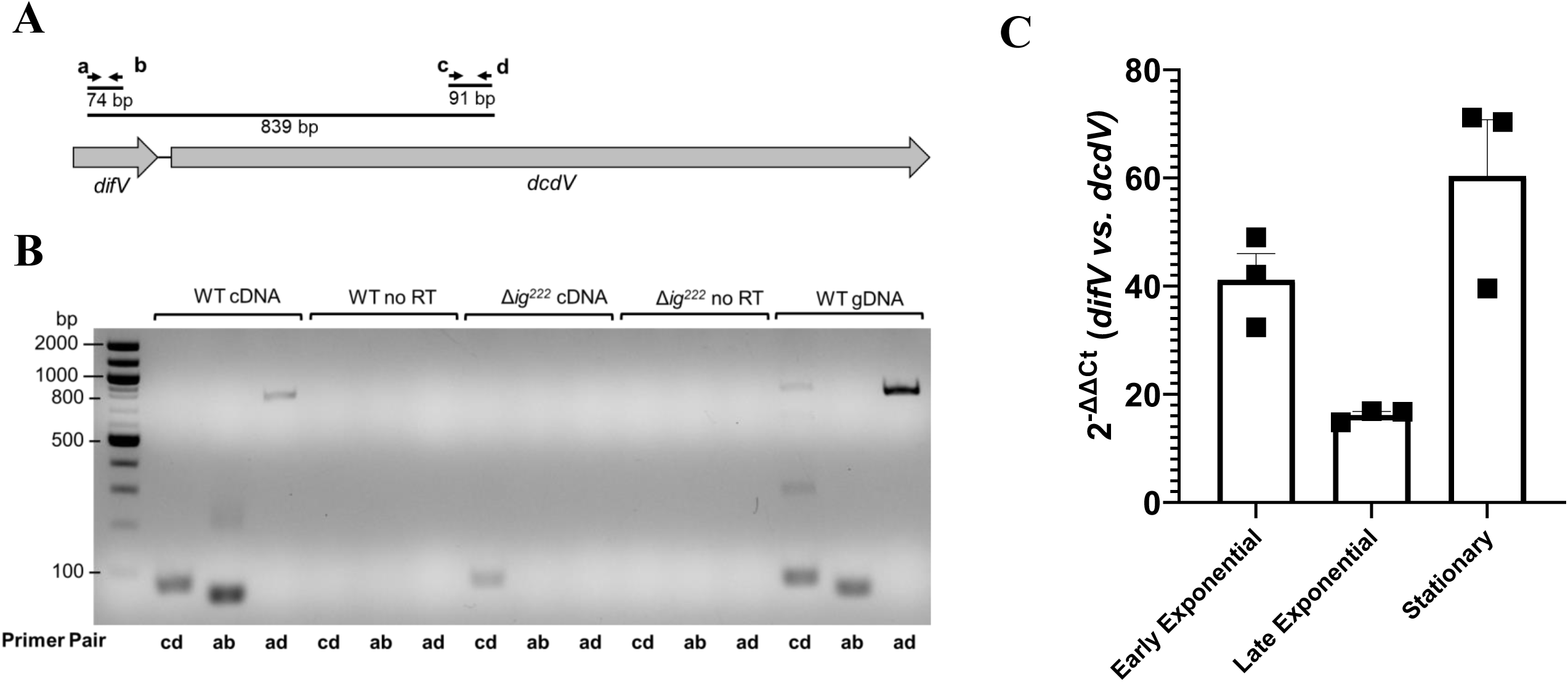
*difV* and *dcdV* are in an operon and *difV* expression exceeds *dcdV*. (**A**) Genomic diagram of *difV* and *dcdV* and the primers (a, b, c, and d) used for generating diagnostic PCR products. (**B**) PCR products amplified from nucleic acid templates (above) using the indicated primer pairs (below) resolved in a 1% agarose gel. All reactions were performed in duplicate using biologically independent samples with similar results. No RT = non-reverse transcribed RNA control. gDNA = genomic DNA control (**C**) qRT-PCR analysis of relative difference between *difV* transcript and *dcdV* transcript levels at different growth phases in WT *V. cholerae* normalized to an endogenous *gyrA* control. Data are graphed as mean ± SEM, *n=* 3.

### DcdV induced filamentation requires conserved features of both the PLK and the CDA domains

DcdV is a 532 amino acid polypeptide composed of two putative domains: an unannotated N-terminal domain and a DCD-like C-terminus (Figs. 4A, 4B). Analysis of the N-terminal domain using Pfam did not reveal any conserved domains. However, Phyre2 [39] and PSI-BLAST searches combined with InterProScan [40, 41] analyses revealed that the N-terminus contained features of the P-loop containing nucleoside triphosphate hydrolase (IPR ID: IPR027417) *aka* P-loop kinase (PLK) enzyme family (Figs. 4A,B and S6). PLKs catalyze the reversible phosphotransfer of the γ-phosphate from a nucleotide triphosphate donor to a diverse group of substrates, depending on the enzyme class, including deoxynucleotide monophosphates. Three structural features commonly found in these enzymes include a P-loop/Walker A motif {GxxxxGK[ST]}, a two helical LID module that stabilizes the donor nucleotide triphosphates, and a Walker B motif {hhhh[D/E], where “h” represents a hydrophobic residue} that is partly involved in coordinating Mg^2+^ [42, 43]. Interrogation of the Phyre2 DcdV model (Fig. 4A), InterProScan predictions, and PSI-BLAST primary sequence alignments (Fig. S6) revealed these three features are likely present in the N-terminal domain, suggesting the N-terminus of DcdV is a PLK domain involved in binding nucleotide substrates and performing a phosphotransfer reaction. The C-terminal DCD domain contains a highly conserved zinc-dependent CDA active site motif ([HAE]X_28_[PCXXC]) (Figs. 4A,B and S6). The constellation of residues that make up the Zn^2+^ binding pocket is composed of three critical amino acids in DcdV; H382, C411, and C414. Zn^2+^ is required for the catalytic deprotonation of water by a conserved glutamate residue (E384 in DcdV) for the hydrolytic deamination of a cytosine base to uridine.

**Fig. 4:**
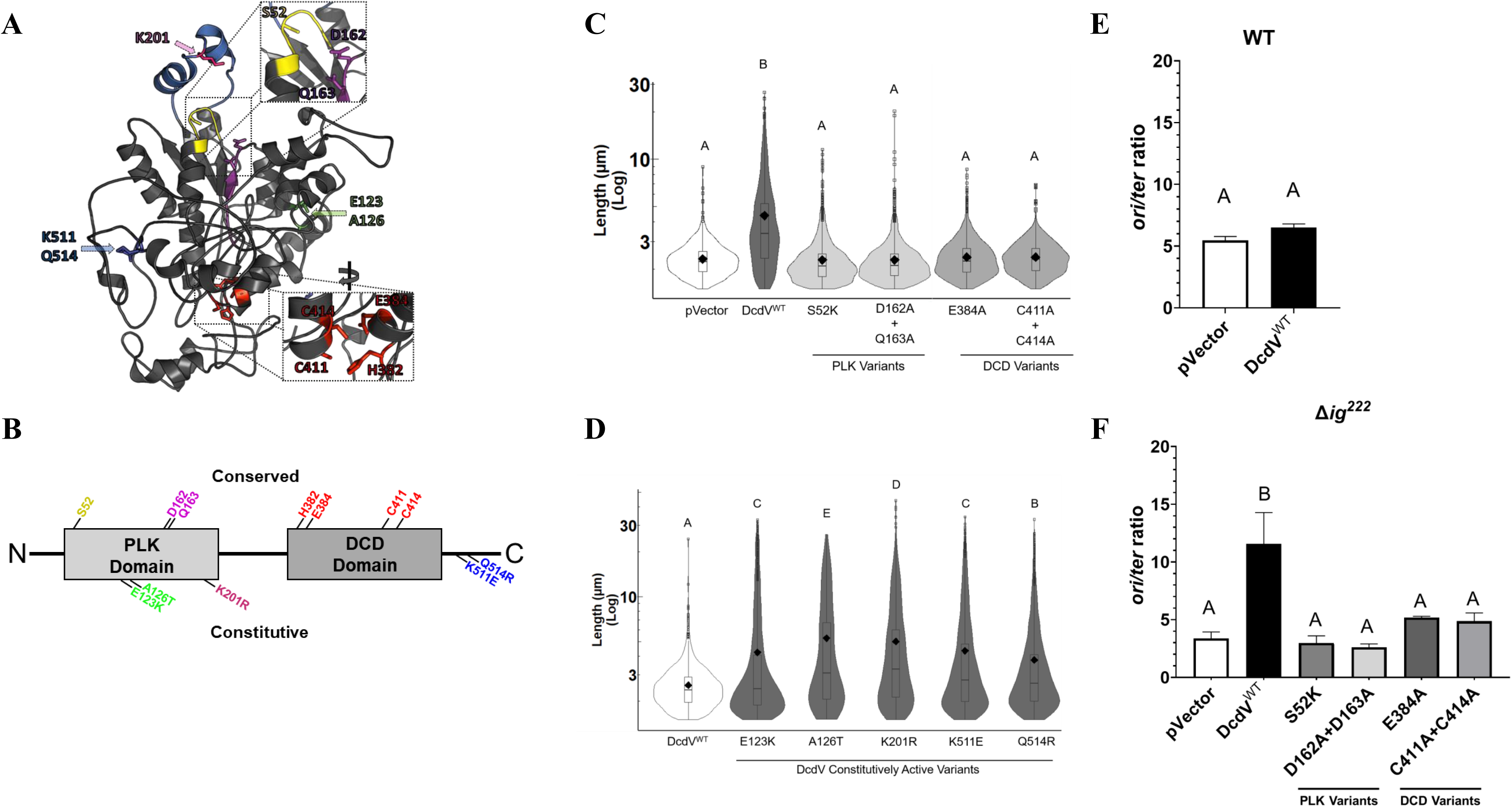
Both the PLK and DCD domains are required for DcdV induced filamentation. (**A**) Phyre2 predicted structure of DcdV from *V. cholerae* El Tor. The inset shows the conserved residues of PLK (top) and DCD (bottom) domains. (**B**) Domain organization and conserved residues at each domain of DcdV. Top labeled residues indicate conserved features of both domains, and the bottom labeled residues indicate variants that render DcdV constitutively active. (**C**) Distribution of cell lengths measured from three biological replicates of WT *E. coli* as indicated. (**D**) Distribution of cell lengths measured from three biological replicates of the Δ*dcdV V. cholerae* mutant expressing the indicated DcdV variants. *ori/ter* ratios of Chromosome 1 in (**E**) WT and (**F**) Δ*ig^222^ V. cholerae* strains expressing the indicated DcdV construct for 8 h and quantified using qRT-PCR. Each bar represents the mean ± SEM, *n*=3. Different letters indicate significant differences (*n*=3) at p < 0.05, according to Tukey’s post-hoc test.

Hypothesizing that one of the two domains present in DcdV is responsible for cell filamentation in the absence of *difV,* we made site-specific mutations in the conserved residues predicted to be essential for activity in both the PLK and DCD domains. Two variant constructs were generated in the PLK domain targeting the Walker A motif (DcdV^S52K^) and the Walker B motif (DcdV^D162A + Q163A^) (Fig. 4B). Two variants were constructed in the DCD active site; a double substitution of both C411A and C414A (DcdV^C411A + C414A^) to abrogate Zn^2+^ binding and an E384A substitution (DcdV^E384A^) to inhibit the deprotonation of water required for the hydrolytic deamination of cytosine (Fig. 4B). Unlike WT DcdV (DcdV^WT^), all four of the variants failed to induce filamentation when ectopically expressed in *E. coli* (Fig. 4C). The cellular abundance of these variants is comparable to WT DcdV (Fig. S7). This result shows both DcdV domains are necessary for induction of filamentation.

We performed a genetic screen to identify DcdV variants whose activity was no longer inhibited by DifV by expressing a random library of *dcdV* mutants in a Δ*dcdV* mutant strain where *difV* remains intact. Ectopic expression of WT *dcdV* in a Δ*dcdV* mutant does not induce filamentation (Fig. 4D) or produce small, wrinkled colonies on solid agar due to the genomic copy of *difV*. However, *dcdV* mutants that are insensitive to *difV* exhibit a small colony phenotype. Screening ∼ 15,000 potential mutants, we identified five unique *dcdV* mutations that encoded single amino acid substitutions (E123K, A126T, K201R, K511E, and Q514R) located in both the PLD and DCD domains that rendered DcdV insensitive to DifV inhibition (Figs. 4B, 4D). Based on the Phyre2 DcdV structural model, all five residues are located on the exterior of the protein (Fig. 4A) suggesting they may be involved in mediating molecular interactions between DifV and DcdV. The only mutation found within a conserved domain feature was the seemingly innocuous K201R substitution, which is modeled to lie between the two helices of the PLK LID module (Fig. 4A).

### DcdV induced filamentation is due to impaired genome replication

Filamentation is a phenotype often associated with thymineless death (TLD) [28] due to nucleotide starvation. A hallmark of TLD is an increased genomic origin to terminus (*ori*/*ter*) ratio resulting from repeated attempts to initiate replication from the origin that ultimately fail to reach the terminus due to a lack of dTTP substrate [44]. Hypothesizing that DcdV induced filamentation may be a consequence of replication inefficiency, analogous to TLD, we measured the *ori/ter* ratio of *V. cholerae* chromosome 1 from WT and Δ*ig^222^ V. cholerae* grown to stationary phase overexpressing WT DcdV or a vector control. There was no significant difference in the *ori/ter* ratios following ectopic expression of WT DcdV in WT *V. cholerae* (Fig. 4E), consistent with the observation that these strains do not filament (Figs. 1D and 1E). However, ectopic expression of WT DcdV in the Δ*ig^222^* mutant, which lacks *difV*, resulted in an *ori*/*ter* ratio ∼ 3 times greater than the vector control (Fig. 4F), consistent with cell filamentation (Fig. 2B). We also measured the *ori*/*ter* ratio of the Δ*ig^222^* mutant expressing *dcdV* with mutations in the PLK or DCD domain. In agreement with an inability to induce filamentation (Fig. 4C), the *ori*/*ter* ratio of these variants was not significantly different from the empty vector control (Fig. 4F). Therefore, DcdV corruption of DNA replication is dependent upon both the PLK and DCD domains.

### DcdV catalyzes the deamination of both dCMP and dCTP

Based on the TLD-like genome instability driven by DcdV, we hypothesized this enzyme deaminates free nucleic acid substrates. Though we determined DcdV and DcdV variants were readily retained in *E. coli* lysates (Fig. S7), numerous attempts to purify active DcdV were unsuccessful. This suggested that an unknown cofactor or cellular condition may contribute to the activity of DcdV that was missing in our purification conditions. Soluble lysates from *E. coli* ectopically expressing DcdV or the DCD active site variant DcdV^E384A^ were supplemented with amine containing nucleotides and monitored for the evolution of NH_4_, a product of nucleotide deamination. Lysates containing DcdV evolved significantly more ammonium when incubated with dCMP and dCTP, which was not detected in lysates containing the DCD active site variant DcdV^E384A^ (Fig. 5A).

**Fig. 5:**
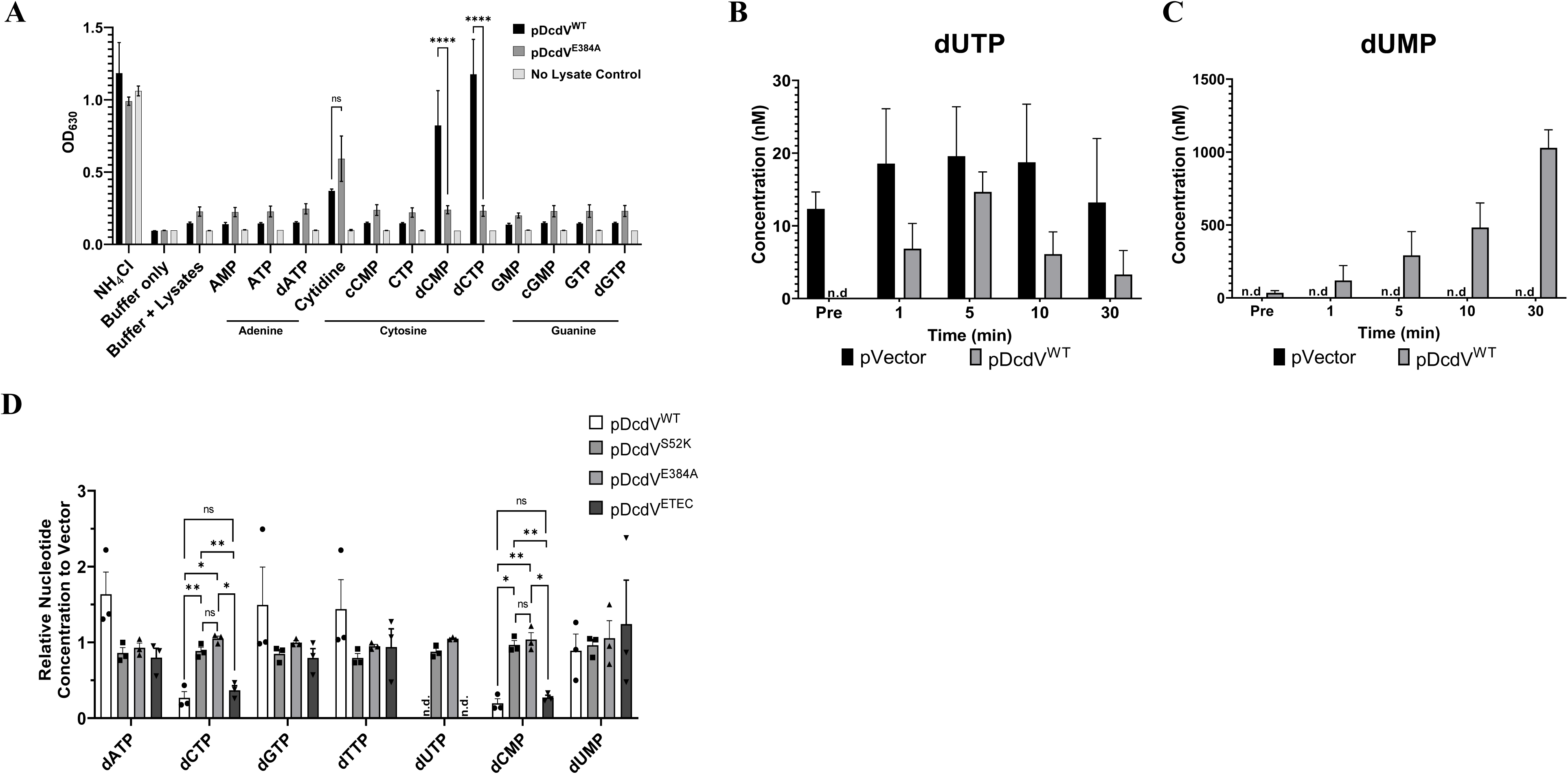
DcdV alters cellular nucleotide metabolism. (**A**) Lysates collected from *E. coli* expressing DcdV or DcdV^E384A^ and a “no lysate” buffer control incubated with 12 nucleotide substrates (1.9 mM NH_4_Cl as a positive control, 37.7 mM cytidine, and 7.5 mM for all other substrates). Data represent the mean ± SEM, *n*=3. Quantification of dUTP (**B**) and dUMP (**C**) using UPLC-MS/MS, in the indicated cell lysates before (Pre) and after addition of 1 mM dCTP. Each lysate was normalized to 20 mg/mL total protein. Each bar represents mean ± SEM, *n*=3. (**D**) Quantification of the indicated dNTPs in vivo using UPLC-MS/MS in strains expressing the four DcdV variants, as indicated, normalized to dNTP concentrations measured in a vector control. Data are graphed as mean ± SEM, *n=* 3, Two-way ANOVA with Tukey’s multiple-comparison test, normalized to pVector, n.d. indicates “none detected”, and ns indicates “not significant”.

DCD enzymes are unique among the CDAs for their allosteric regulation by both dCTP and dTTP which activate and repress the catalytic deamination of dCMP, respectively, through a G[Y/W]NG allosteric site motif [45, 46]. Such allosteric regulation ensures that nucleotide homeostasis is maintained even if DCD enzymes are present. The allosteric site found in DcdV is composed of a divergent GCND motif suggesting allosteric regulation by dNTPs may not be preserved. In support of this, the deamination of both dCMP and dCTP by soluble lysates containing DcdV were not inhibited by the addition of equimolar dTTP (Fig. S8).

To further understand the catalytic activity of DcdV we spiked 1 µM dCTP into soluble lysates collected from *E. coli* ectopically expressing either WT DcdV or a vector control and quantified the concentrations of dUTP and dUMP over 30 minutes using UPLC-MS/MS. Following addition of 1 µM dCTP the concentrations of both dUTP (Fig. 5B) and dUMP (Fig. 5C) increased in lysates containing DcdV within the first minute while those found in vector control lysates did not dramatically change over the course of the entire experiment. The concentration of dUTP in DcdV containing lysates peaked after five minutes and slowly receded over time (Fig. 5B) while the concentration of dUMP in these lysates continued to increase to a final concentration of ∼ 1 µM after 30 minutes (Fig. 5C). Importantly, the equimolar stoichiometry of the1 µM dCTP substrate spike and the 1 µM dUMP detected at the conclusion of the experiment demonstrates that DcdV does not modify nucleotides in a unique manner which would alter their mass. Together these experiments indicate that DcdV deaminates both dCTP and dCMP substrates and DcdV containing lysates ultimately funnel dCTP to dUMP, indicating DcdV is likely to have profound effects on intracellular nucleotide metabolism.

### DcdV decreases intracellular dCTP, dCMP, and dUTP in *E. coli*

Our genetic and in vitro evidence suggested that DcdV catalyzes the deamination of both dCMP and dCTP to the detriment of DNA replication. To quantify the impact of DcdV activity on the intracellular concentrations of deoxyribonucleotide species, we overproduced DcdV, DcdV^S52K^, DcdV^E384A^, and an empty vector control in *E. coli* and measured the abundance of these molecules by UPLC-MS/MS. While all strains contained similar levels of dATP, dGTP, dTTP, and dUMP, the intracellular abundance of dCTP and dCMP were significantly reduced in *E. coli* expressing WT DcdV (Figs. 5D, S9). No dUTP was found following expression of WT DcdV while trace amounts of dUTP were detected in the vector and the two DcdV variant strains (Figs. 5D, S9). Unlike the results observed with the in vitro DcdV lysates (Fig. 5C), no increase in intracellular dUMP concentrations were observed when DcdV was expressed. We speculate the difference between dUMP detected in lysate versus in vivo extractions are due to compensatory metabolic pathways active in live cells which are lost in the lysates. Similar results were obtained when a DcdV homolog derived from enterotoxigenic *E. coli* (DcdV^ETEC^), discussed later in this study, was overexpressed in the same heterologous *E. coli* host (Figs. 5D, S9). Importantly, inactivating amino acid substitutions in conserved features of the PLK (DcdV^S52K^) or DCD (DcdV^E384A^) domains blocked DcdV activity, indicating both domains are necessary for the DcdV dependent depletion of intracellular dC pools (Figs. 5D, S9).

### Conservation and evolution of DcdV

To identify if DcdV is widely conserved, we used six DcdV homologs as starting points from *V. cholerae*, *Vibrio parahaemolyticus*, *E. coli*, *Proteus mirabilis*, *Aeromonas veronii*, and *Enterobacter cloacae* to perform homology searches across the tree of life (see Methods). We used a combination of protein domain and orthology databases, homology searches, and multiple sequence alignment for detecting domains, signal peptides, and transmembrane regions to reconstruct the domain architectures of the query DcdV proteins (Fig. S6). In agreement with the Phyre2 model of *V. cholerae* DcdV (Fig. 4A), we identified two distinct domains in all six DcdV homologs, the N-terminal PLK domain and the C-terminal DCD domain (Fig. S6).

We identified numerous homologs containing the core PLK+DCD architecture as well as other variations, which included multiple PLK domain fusions in proteobacteria (*e.g., Klebsiella*, *Vibrio*) and a nucleic acid binding domain (*e.g., Mannheimia*, *Bibersteinia*) (Table S4). Homologs of DcdV were identified in multiple bacterial phyla including Proteobacteria, Actinobacteria, Bacteroidetes, and Firmicutes (Figs. 6A, a few dominant clusters of homologs are labeled). Interestingly, we found DcdV-like proteins in Archaea (*e.g.,* Thaumarchaeota) and Eukaryota (*e.g.,* Ascomycota) (Figs. 6A, Table S4). While the percentage similarity is ∼50% and <30% for archaeal and eukaryotic homologs, respectively, we note these contain comparable domain architectures to the query proteins (Table S4).

### Identification and evaluation of Gram-negative DcdV-DifV system homologs

To evaluate the conservation of enzymatic activity we selected three of the core DcdV homologs used in the initial homolog search; *V. parahaemolyticus* O1:Kuk FDA_R31*, P. mirabilis* AR379, and *E. coli* H10407 ETEC (Figs. 6A and S10). Expression of all three DcdV homologs in *E. coli* resulted in filamentous cells analogous to *V. cholerae* DcdV (Fig. 6B). These *dcdV* homologs are encoded 3’ of a small ORF, annotated as a hypothetical protein, in an orientation, size, and proximity consistent with *V. cholerae difV*. While there was no strong amino acid or nucleotide sequence similarity among the small ORFs 5’ of the *dcdV* homologs (Figs. S11 and S12) we hypothesized these could encode cognate *difV* negative regulators. Consistent with the inhibition of DcdV activity by DifV from *V. cholerae*, co-expression of the corresponding DifV with its DcdV partner suppressed the cell filamentation phenotype (Fig. 6B). Additionally, overexpression of DcdV^ETEC^ in a heterologous *E. coli* host also decreased the intracellular concentrations of dCMP, dCTP, and dUTP (Fig. 5D), indicating the catalytic activity of these DcdV homologs are analogous to *V. cholerae* DcdV.

**Fig. 6:**
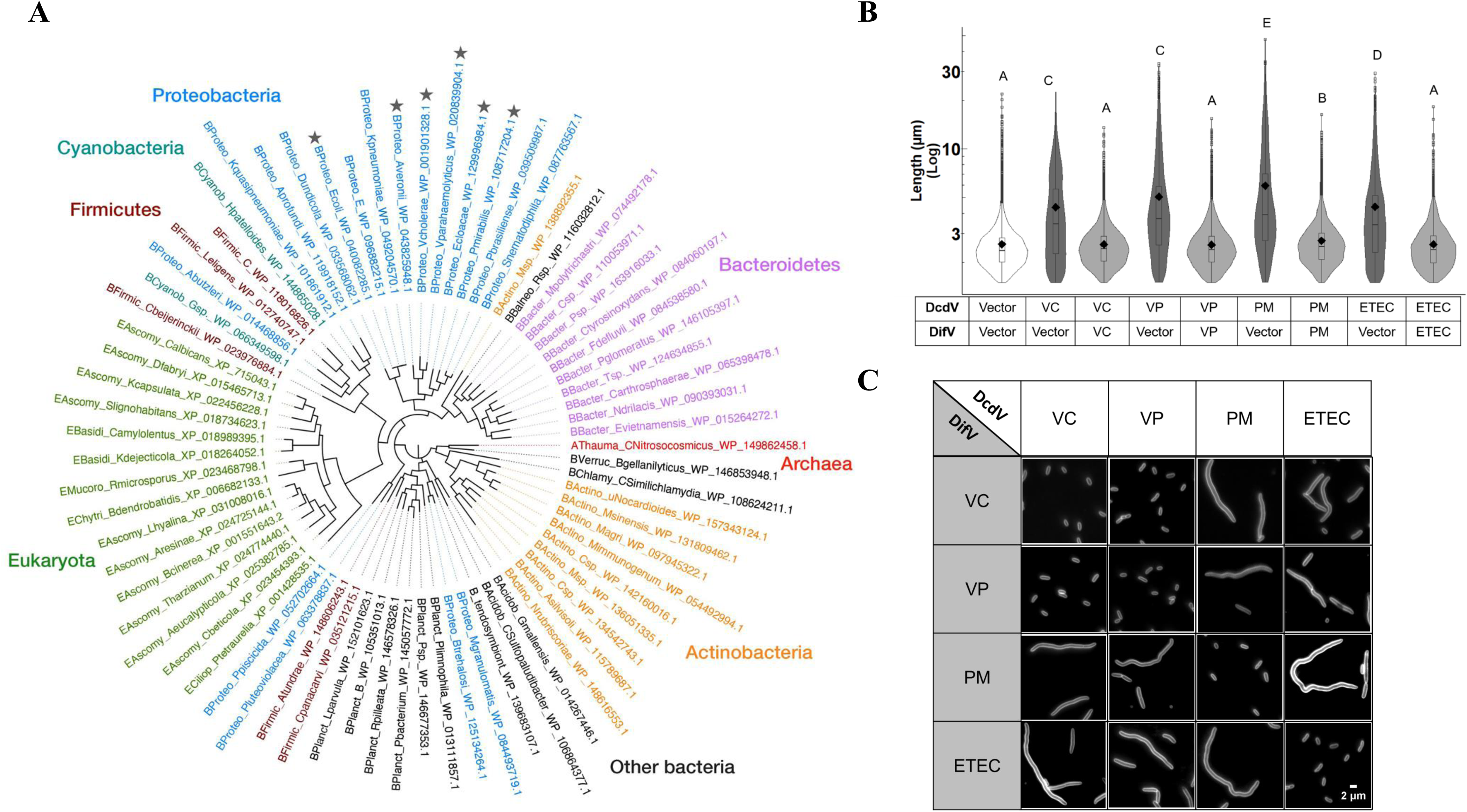
*dcdV* and *difV* are widely conserved. (**A**) Phylogenetic tree of DcdV homologs containing PLK and DCD domains from representative phyla across the tree of life. Stars indicate query proteins of interest in this study. (**B**) Distribution of cell lengths expressing the indicated DcdV homologs and their cognate DifV or vector control in *E. coli* (n=3). Different letters indicate significant differences (*n*=3) at *p*< 0.05, according to Dunnett’s post-hoc test against the control (pVector^DcdV^ + pVector^DifV^) strain. (**C**) Representative images of *E. coli* expressing pDcdV/homologs and pDifV/homologs combinations. Scale represents 2 µm.

To determine if DifV and the three ORFs encoded upstream of *dcdV* homologs could provide cross-species inhibition of DcdV, we challenged each of the four *dcdVs* with each of the four *difVs* in *E. coli* and looked for DcdV dependent filamentation. Cross-species inhibition of DcdV induced filamentation was observed between *V. parahaemolyticus* and *V. cholerae* when each species’ *difV* was expressed in trans (Fig. 6C). However, *difV* from *P. mirabilis* and *E. coli* ETEC were only able to inhibit the activity of their own cognate DcdV (Fig. 6C). These data suggest that while the general mechanism of DifV inhibition of DcdV activity is conserved the specific molecular interactions that mediate this process are not.

### Ectopic expression of DcdV reduces phage titers and slows predation

We initiated studies of *dcdV* based on our discovery that this gene co-occurs in bacterial genomes with *dncV*, a critical member of the CBASS antiphage abortive infection system [10, 47]. Additionally, cytidine deaminases are conserved anti-viral defense mechanisms in eukaryotes [15, 17, 48]. These connections led us to hypothesize that DcdV can also provide phage defense by manipulating cellular nucleotide concentrations. To test this hypothesis, we challenged *V. cholerae* WT and Δ*dcdV* with two *V. cholerae* lytic phage with dsDNA genomes, ICP1 and ICP3 [49, 50]. However, we observed no differences in the ability of these phages to kill *V. cholerae* in these conditions (Figs S13A and S13B).

Because ICP1 and ICP3 have coevolved with El Tor *V. cholerae,* it is likely that these phages have evolved mechanisms to counteract *dcdV*. Such resistance to other *V. cholerae* phage defense mechanisms by ICP-1 has been previously demonstrated [51–53]. Therefore, we selected the heterologous host *Shigella flexneri,* a Gram-negative human pathogen, and its bacteriophage Sf6, a dsDNA phage from the *Podoviridae* family [54, 55], as a naïve host-phage pair to test the antiphage activity of DcdV and its homologs. Ectopic expression of *dcdV* or its homologs did not impact the growth of *S. flexneri* before the onset of phage killing at ∼110 minutes (Figs. 7A-D). *S. flexneri* strains ectopically expressing *dcdV* or its homologs delayed the onset of population collapse caused by Sf6 predation, although the impact of the *V. cholerae* DcdV was more modest than the other three homologs (Figs. 7A-D). Additionally, induction of all four DcdV homologs significantly reduced Sf6 progeny following infection compared to the control strains lacking induction of DcdV (Fig. 7E). Together, these data indicate that DcdV enzymes confer defense against phage infection by delaying population collapse and reducing the proliferation of viable phage progeny.

**Fig. 7:**
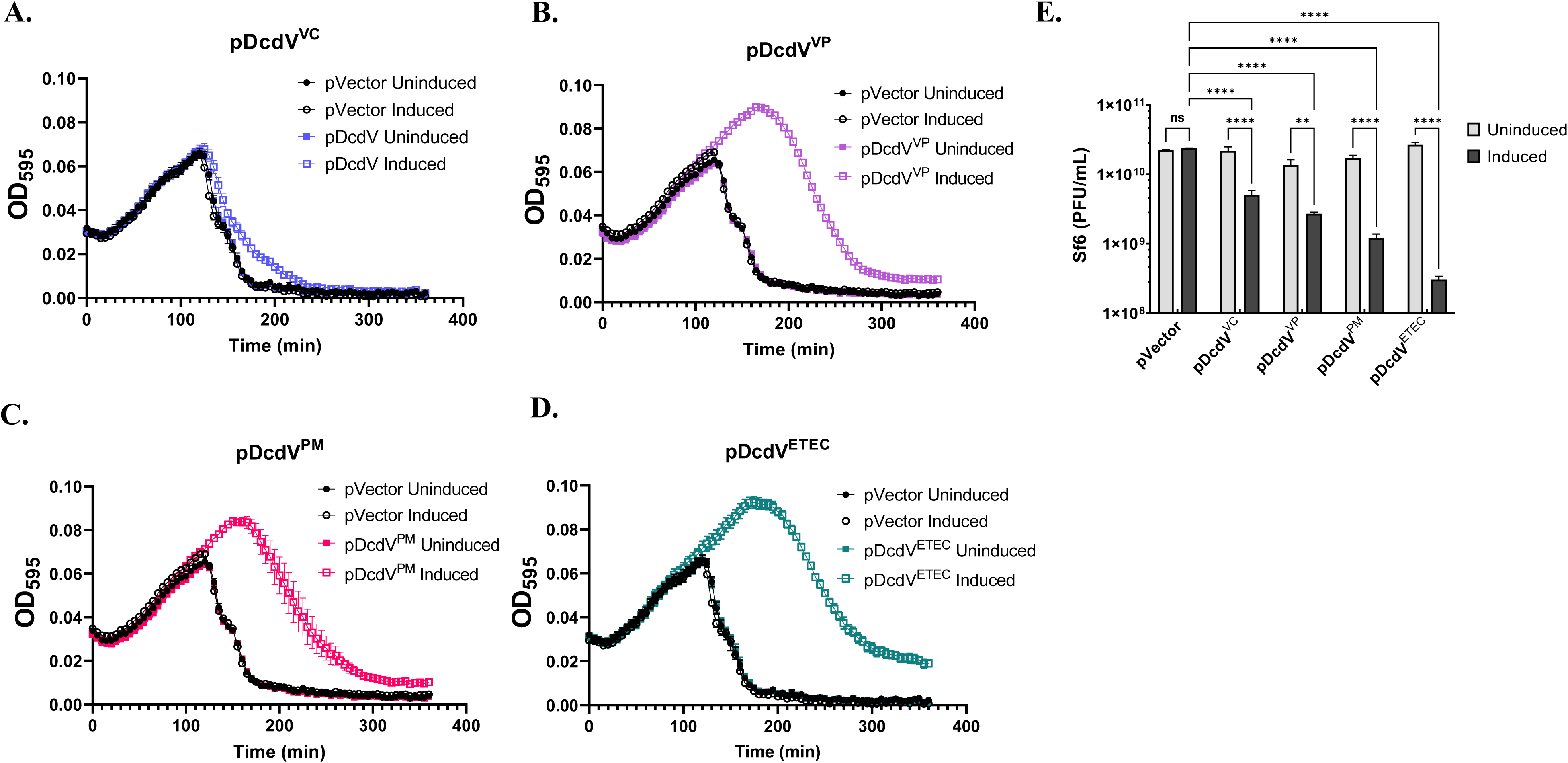
DcdV mediates phage defense. (**A-D)** Growth curves for *S. flexneri* containing vector or pDcdV/homologs infected Sf6 at time 0 at an MOI of 0.1 in the presence or absence of 100 µM IPTG. Each graph represents three biological replicates each with three technical replicates. (**B**) Plaque-forming units (PFU) per mL of phage Sf6 measured at the conclusion of the *S. flexneri* growth curve experiment above. Results are represented as mean ± SEM, *n=* 3, Two-way ANOVA with Tukey’s multiple-comparison test.

## DISCUSSION

Uncovering the contributions to bacterial fitness of the ∼36 genes encoded within the El Tor *V. cholerae* VSP-1 and 2 genomic islands may help elucidate the longevity and persistence of the seventh cholera pandemic. Our bioinformatic approach using Correlogy accurately identified a gene network composed of the VSP-1 antiphage CBASS system (*capV-dncV-vc0180-vc0181*). Interestingly, this also revealed *dncV* is frequently found in genomes with the previously uncharacterized gene *dcdV*. The only function previously ascribed to *dcdV* was an undefined involvement in quorum sensing controlled *V. cholerae* aggregate formation [56].

We showed that DcdV contains a functional DCD domain that catalyzes the deamination of deoxycytidine nucleotides and a putative PLK-like domain of unknown function. We further demonstrate that homologs of this protein are present across the tree of life. Collectively, both domains are required for DcdV to disrupt deoxynucleotide pool homeostasis, which impairs DNA replication and manifests in a filamentous cell morphology. DcdV activity is post-translationally regulated by DifV, a sRNA encoded immediately 5’ of the *dcdV* locus in VSP-1, though the details of this inhibition remain to be fully elucidated. Finally, we demonstrate that DcdV and a set of homologs from other Gram-negative bacteria confer phage resistant properties when expressed in a heterologous host.

Cell filamentation is a hallmark of TLD, observed in bacteria and eukaryotes, which arises from a sudden loss of thymine during robust cellular growth [31]. Interestingly, this phenomenon is not limited to dTTP as dGTP starvation elicits a similar response in *E. coli* and is also hypothesized to occur when other deoxynucleotide substrates become disproportionately scarce [29]. In the case of DcdV, it is conceivable the observed filamentation phenotype is a consequence of a TLD-like reduction in dCTP pools that can be termed ‘*cytosineless death’*.

However, while DcdV activity also reduces the intracellular dC pool, it did not significantly increase the intracellular concentrations of dTTP or dUMP in vivo, suggesting a cellular compensatory pathway to combat DcdV activity is at work in intact cells. We speculate that the DCD and PLK domains of DcdV are responsible for this conversion of dC nucleotides to dUMP observed in the bacterial lysates, but we cannot rule out the contribution of other unknown cellular factors. The deamination of dCTP is canonically performed by non-zinc dependent enzymes [57] making the dual substrate repertoire of dCMP and dCTP in DcdV a rare trait.

The delicate balance of enzymatic activity across the pyrimidine biosynthesis pathway can be corrupted by viruses that deploy their own DCD, dUTPase, and TS enzymes to hijack host nucleotide biosynthesis to ensure the appropriate ratio and quantities of deoxyribonucleotide precursors for replicating their own genomes [23, 24, 26, 27]. For example, biDCD from chlorovirus PBCV-1, the only DCD previously reported to deaminate both dCMP and dCTP substrates, rapidly catalyzes the conversion of host dC nucleic acids into dTTP thus aiding replication of the A+T rich viral genome [23]. biDCD is allosterically regulated by dCTP and dTTP to activate and inactivate the deaminase, respectively. This regulation provides a means to fine-tune the pool of available dNTPs by preventing the enzyme from deaminating all available dC substrates. Interestingly, DcdV does not appear to have maintained the allosteric nucleotide binding site nor does excess dTTP added to cell lysates alter the catalytic activity of DcdV towards dCMP or dCTP (Fig. S8), and we propose these differences in enzyme activity are consistent with the function of DcdV as a phage defense mechanism that inhibits phage replication by corrupting cellular nucleotide pools (graphical abstract). Altering pools of available nucleotides has been shown to fend off biological attacks. For example, prokaryotic viperins protect against T7 phage infection by producing modified ribonucleotides that ultimately inhibit phage polymerase-dependent transcription [58]. The SAMHD1 phosphohydrolase enzyme in eukaryotes also inhibits viral infections by depleting cellular nucleotide pools, although its structure and activity are different than DcdV [59–61].

In lieu of a conserved deoxynucleotide allosteric site, DcdV is regulated post-translationally by the DifV untranslated RNA, which is unique among the CDA-family. The spacing, orientation, and relationship of *difV* and *dcdV* may have adapted to perform functions in a manner analogous to Type 2 and Type 3 Toxin-Antitoxin (TA) systems found across the bacterial phyla of which some are involved in antiphage defense and bacterial stress response [62]. While the RNA antitoxin of Type 3 TA systems encode nucleotide repeats [62] no repeat sequences are obvious in DifV indicating that DcdV/DifV may constitute a new TA class. We hypothesize that DcdV is activated upon phage infection by disruption of DifV inhibition, and we are currently preforming experiments to test this hypothesis (graphical abstract). Our systemic search for DcdV homologs containing at least a single PLK and DCD domain revealed hundreds of examples in a variety of bacteria beyond the Proteobacteria phylum including Bacteroidetes and Actinobacteria and a few homologs in archaea and eukaryota.

Phage defense mechanisms are often found clustered together in mobile genetic elements called defense islands [63, 64] and we speculate that the co-occurrence of DcdV and DncV (along with the rest of the CBASS system) in bacterial genomes is a result of their shared anti-phage activity. Our results indicate that DcdV reduces the available dC pool, and we hypothesize that this activity delays phage genome replication potentially decreasing phage burst size. Although the *S. flexneri* host population expressing DcdV eventually collapses, we speculate that the delay in phage replication could provide an opportunity to prompt other phage defense systems, such as CBASS or a restriction modifications system to further target invading phages [65, 66].

Our study reveals that bacteria, like eukaryotes, also use CDA enzymes to protect against biological invasion although through different mechanisms. The eukaryotic APOBEC proteins deaminate ssRNA, leading to increased mutation and decreased genome stability of RNA viruses, whereas the substrates of DcdV are free deoxynucleotides. Further studies are required to determine if these two biological defense systems evolved from a common CDA ancestor.

## ACKNOWLEDGEMENTS

We thank Shannon Manning (STEC Center, Michigan State University), Jessica Jones (U.S FDA), and Allison Brown (U.S. CDC), for providing us with *E. coli* ETEC, *P. mirabilis,* and *V. parahaemolyticus* strains, respectively. We thank Wei Leung Ng (Tuft University) for providing us *V. cholerae* ICP1 and ICP3 phages. We thank Kefei Yu and Dohun Pyeon for valuable suggestions and Dan Jones and Lijun Chen from the MSU RTSF mass spectrometry facility core for their technical support. This work was supported by National Institutes of Health (NIH) grants GM109259, GM110444, GM139537, AI143098, and AI158433 and National Science Foundation (NSF) grant DBI-0939454to C.M.W, the NIH grant GM110185 and the NSF CAREER Award 1750125 to K.N.P., and the NSF Graduate Research Fellowship Grant No. 1842399 to C.A.E. Any opinions, findings, and conclusions or recommendations expressed in this material are those of the author(s) and do not necessarily reflect the views of the National Science Foundation.

## MATERIALS AND METHODS

The strains, plasmids, and primers used in this study are listed in Supplementary Table S1, S2, and S3, respectively. Unless otherwise stated, cultures were grown in Luria-Bertani (LB) at 35°C and supplemented with the following as needed: ampicillin (100 µg/mL), kanamycin (100 µg/mL), tetracycline (10 µg/mL), and isopropyl-β-D-thiogalactoside (IPTG) (100 µg/mL). *E. coli* BW29427, a diaminopimelic acid (DAP) auxotroph, was additionally supplemented with 300 μg/mL DAP. The *V. cholerae* El Tor biotype strain C6706str2 was utilized in this study and mutant strains were generated using the pKAS32 suicide vector [67] using three fragments: 500 bp of sequence upstream of the gene of interest, 500 bp of sequence downstream of the gene of interest and cloned into the KpnI and SacI restriction sites of pKAS32 using by Gibson Assembly (NEB). P_tac_ inducible expression vectors were constructed by Gibson Assembly with inserts amplified by PCR and pEVS143 [68] or pMMB67EH [69] each linearized by EcoRI and BamHI, as well as pET28b digested with NcoI and XhoI. pEVS141 [70] is used as an empty vector control for experiments using pEVS143 derived constructs. Site-directed mutagenesis was performed using the SPRINP method [71]. Plasmids were introduced into *V. cholerae* through biparental conjugation using an *E. coli* BW29427 donor. Transformation of *E. coli* for ectopic expression experiments was performed using electroporation with DH10b for expression of pEVS143 and pMMB67EH derived plasmids and BL21(DE3) for pET28b based constructs.

### Correlogy Bioinformatics Analysis

Our Correlogy software package is built on Kim and Price’s approach [32] to calculate genetic co-occurrence. The source code, documentation, and a Docker container for this Python3 package are available at https://github.com/clinte14/correlogy. While VSP-1 is used to simplify the description of the method detailed below, both VSP-1 and 2 were independently analyzed in the same fashion. To establish maximum related subnetworks (MRS) for the genomic region of the VSP-1 island, a BLASTP amino acid sequence was performed to search for each VSP-1 gene against the NCBI non-redundant protein database with an E-value cutoff of 10^-4^. The BLAST results were limited to bacterial genomes, and all taxa belonging to the genus *Vibrio* were removed to avoid bias from closely related vertical inheritance. The BLAST results were used to generate a presence or absence matrix of VSP-1 homologues with all species along one axis and VSP-1 genes along the other axis. Next, a pairwise Pearson correlation value was calculated between all VSP-1 genes *i* and *j* using binary data from the above-mentioned presence/absence matrix:

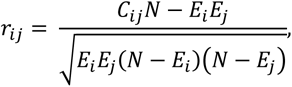

where N is the total number of unique species returned from the BLAST search and *C_ij_* the number of species with co-occurrence of genes *i and j.* While a Pearson correlation is warranted for a normally distributed binary data set, it does not account for indirect correlation. For example, if genes *i* and *j* individually associate with a third gene, a Pearson correlation will incorrectly calculate a correlation between *i* and *j.* To help correct for indirect correlation we calculate a partial correlation *w_ij_* from the Pearson *r_ij_*:

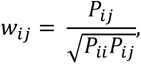

where the (*i, j*) element of the inverse matrix of Pearson *r_ij_* is *P_ij_* [32].

The partial correlation correction *w_ij_* has the advantage of generating a normalized output that ranges between -1 to 1. For example, a *w_ij_* of -1 reveals genes *i* and *j* never occur in the same species, while a value of 1 demonstrates genes *i* and *j* always co-occur in the same species. A *w_ij_* of 0 is the amount of co-occurrence expected between unrelated genes *i* and *j* drawn from a normal distribution. Using the above-mentioned approach, a partial correlation value *w_ij_* was calculated for all genes *i* to *j* in VSP-1 and VSP-2 (Supplemental Files 1 and 2). The single highest *w_ij_* value for each VSP-1 gene was represented as an edge (i.e., line) in our visualization (Fig. 1B, S1A, and S1B). Any set of genes that contains no further edges were assigned to a unique MRS that suggests functional association of the gene products within a unique gene network.

### Genomic Identification, Structural, and Sequence Analyses of DcdV & DifV Homologs

DcdV from *V. cholerae* El Tor N16961 (WP_001901328.1) was identified as locus tag *vc0175*. DcdV and homologs profiles are performed using translated BLAST tblastn and run against the Nucleotide collection (nr/nt) database of National Center for Biotechnology Information (NCBI), using >40% similarities cutoff. For previously annotated domains, the Pfam feature in KEGG [72, 73] were utilized as a guide to determine DcdV homologs. Out of all the DcdV homologs, DcdV homologs from *Vibrio parahaemolyticus* O1: Kuk str. FDA_R31 (WP_020839904.1), *Proteus mirabilis* AR_379 (WP_108717204.1), and *E. coli* O78:H11 H10407 (ETEC) (WP_096882215.1) were analyzed in this study. Genomic contextual information from prokaryotic gene neighborhoods was retrieved from NCBI genome graphics feature to uncover *difV*-like gene, encoded as a hypothetical ORF 5’ of the *dcdV* locus. If unannotated, the ORFinder feature from NCBI was used to determine the location and size of the putative *difV* locus. To predict the structure of DcdV from *V. cholerae,* the amino acid sequence was submitted to Phyre2 [39] and structural visualization was performed using PyMol (https://pymol.org). The amino acid and nucleotide alignments were analyzed using ClustalW Omega from EMBL-EBI web services [74] and LocARNA [75], respectively.

### Identification and Characterization of Protein Homologs

**Homology searches**: To ensure the identification of a comprehensive set of homologs (close and remote), we started with six representative DcdV proteins across proteobacteria from *V. cholerae*, *V. parahaemolyticus*, *P. mirabilis*, and *E. coli* described above along with *E. cloacae* (WP_129996984.1), and *A. veronii* (WP_043825948.1) and performed homolog searches using DELTABLAST [76] against all sequenced genomes across the tree of life in the NCBI RefSeq database [77–79]. Homology searches were conducted for each protein and the search results were aggregated; the numbers of homologs per species and of genomes carrying each of the query proteins were recorded. These proteins were clustered into orthologous families using the similarity-based clustering program BLASTCLUST [76].

**Characterizing homologous proteins**: Phyre2, InterProScan, HHPred, SignalP, TMHMM, Phobius, Pfam, and custom profile databases [39–41, 80–85] were used to identify signal peptides, transmembrane (TM) regions, known domains, and secondary structures of proteins in every genome. Custom scripts were written to consolidate the results [86–91], and the domain architectures and protein function predictions were visualized using the MolEvolvR web-app (http://jravilab.org/molevolvr/).

**Phylogenetic analysis (MSA and Tree)**: Thousands of homologs from all six starting points for DcdV proteins were consolidated and representatives were chosen from distinct Lineages and Genera, containing both the N- and C-terminal DcdV domains (PLK and DCD domains). Multiple sequence alignment (MSA) of the identified homologs was performed using Kalign [89] and MUSCLE [92, 93] (msa R package [94]). The phylogenetic trees were constructed using FastTree [95] FigTree [96] and the R package, ape [97].

### Growth Curve Assays

Overnight cultures were diluted 1:1000 into LB supplemented with antibiotics and IPTG in a 96-well microplate (Costar®). Growth was monitored by measuring OD_600_ every 15 minutes for 15 hour (h) using a BioTek plate reader with continuous, linear shaking.

### Fluorescence Microscopy and Analysis

Cells were imaged as previously described [34]. Briefly, overnight cultures were diluted 1:1000 into LB supplemented with antibiotics and IPTG. Cultures were grown and induced for 7-8 h, at which point cells were diluted to an OD_600_ of 0.5 in 1X PBS, then membrane stain FM4-64 dye (ThermoFisher Scientific) was added to a final concentration of 20 µg/mL. 1% agarose pads in deionized water were cut into squares of approximately 20 x 20 mm and placed on microscope slides. 2 µl of diluted cultures were spotted onto a glass coverslip and then gently placed onto the agarose pad. FM4-64 signal was visualized using a Leica DM5000b epifluorescence microscope with a 100X-brightfield objective under RFP fluorescence channel. Images were captured using a Spot Pursuit CCD camera and an X-cite 120 Illumination system. Each slide was imaged with at least 20 fields of view for each biological replicate. Cell lengths were processed using the Fiji plugin MicrobeJ [98, 99], and data were visualized and analyzed using R [90] by quantifying the length of the curvilinear (medial) axis of detected cells.

### Construction and screening of mutant gene libraries

DifV-insensitive DcdV constructs were generated by error-prone PCR (epPCR) using pDcdV (pCMW204) as the template. Three different concentrations of MnCl_2_ (12.5 mM, 1.25 mM, and 125 μM) were used in triplicate using Taq polymerase (Invitrogen) and reactions containing the same MnCl_2_ concentration were pooled. The PCR products were purified, using The Wizard® SV Gel *and* PCR Clean*-*Up Kit (Promega), and ligated to pEVS143 via Gibson Assembly. The assembled reactions were electroporated to *E. coli* DH10b and plasmid libraries were collected from ∼ 30,000 representative colonies for each MnCl_2_ concentration. Plasmid libraries were harvested using the Wizard® Plus SV Minipreps DNA purification Kit (Promega). Plasmid libraries were subsequently electroporated to *E. coli* BW29427 which were again plated and pooled to contain ∼ 30,000 representative colonies. The *E. coli* BW29427 random mutant pDcdV libraries were conjugated with Δ*dcdV V. cholerae* on LB agar plates for 8 h, harvested, diluted, and spread on LB agar plates containing 1 mM IPTG and antibiotics, and grown overnight. ∼ 5,000 colonies were screened in each library and all colonies exhibiting a wrinkled and small colony morphology, indicative of cell filamentation, were isolated and filamentation was confirmed by fluorescence microscopy. Mutant pDcdV plasmids recovered from cells exhibiting cell filamentation were sequenced by Sanger sequencing. Mutations were reintroduced individually into the WT pDcdV construct using SPRINP mutagenesis [71] and reevaluated using fluorescence microscopy to confirm the DcdV variant’s ability to remain constitutively active in Δ*dcdV V. cholerae*.

### RNA Isolation, qRT-PCR, and Co-transcription Analysis

RNA isolation and qRT-PCR analysis were carried out as previously described [100, 101]. Briefly, triplicate overnight cultures were subcultured 1:1000 in 10 mL LB and grown to three different OD_600_: 0.2 (Early Exponential), 1.0 (Late Exponential), and 2.5 (Stationary). 1 mL of each replicate was pelleted, and RNA was extracted using TRIzol^®^ reagent following the manufacturer’s directions (Thermo Fischer Scientific). RNA quality and quantity were determined using a NanoDrop spectrophotometer (Thermo Fischer Scientific). 5 µg of purified RNA was treated with DNase (Turbo^TM^ DNase, Thermo Fischer Scientific). cDNA synthesis was carried out using SuperScript^TM^ III Reverse Transcriptase (Thermo Fischer Scientific). cDNA was diluted 1:64 into molecular biology grade water and amplification was quantified using 2x SYBR Green (Applied Biosystems^TM^). For measuring gene expressions or determining *ori*/*ter* ratios, 25 µL reactions consisted of 5 µL each of 0.625 µM primers 1 and 2, 12.5 µL of 2X SYBR master mix, and 2.5 µL of template (0.78 ng/μL cDNA for gene expression and 0.25 ng/μL genomic DNA for *ori*/*ter*). qRT-PCR reactions were performed in technical duplicates for biological triplicate samples and included no reverse transcriptase reaction controls (“no RT”) to monitor for contaminating genomic DNA in purified RNA samples. qRT-PCR reaction thermo profile was 95°C for 20 seconds (s) then 40 cycles of 95°C for 2 s and 60°C for 30 s in the QuantStudio 3 Real-Time PCR system (Applied Biosystems^TM^). The *gyrA* gene was used as an endogenous control to calculate relative quantification (Δ*C*_t_).

To determine the co-transcription of *difV* and *dcdV*, PCR amplification was performed in 25 µL volumes using Q5 polymerase (NEB), 0.5 µM each of the forward and reverse primers as indicated, 0.2 mM dNTPs, and 3.5 µL of cDNA or no RT control templates (0.78 ng/µL) from RNA purified from WT and Δig^222^ *V. cholerae* grown to late exponential-phase in biological triplicate. The thermal profile was 98°C for 30 s, 30 cycles of 98°C for 10 s, 55 °C for 30 s, 72 °C for 10 sec and one cycle of 72 °C for 2 min. PCR products were loaded on a 1% agarose gel and stained with EZ-Vision® (VWR). Images were taken using the GelDoc system (Bio-Rad).

### In-vitro Nucleic Acid Deamination Assay

**Cell Lysate Preparation:** Overnight cultures were subcultured 1:333 and grown to an OD_600_ of ∼0.5 - 1.0. Cultures were induced with 1 mM IPTG, supplemented with 100 µM ZnSO_4_, and grown for an additional 3 hr. Cell pellets from 100 mL of induced cultures were harvested in two successive 15 min centrifugation steps at 4,000 x g and 4°C. Supernatants were decanted and pellets were snap frozen in an ethanol and dry ice bath and stored at -80° C. Pellets were thawed on ice and suspended in 2 mL of lysis buffer A (50 mM NaPO_4_, pH 7.3, 300 mM NaCl, 2 mM β-mercaptoethanol, 20% glycerol and Roche cOmplete protease inhibitor (1 tablet per 10 mL)). 1 mL of cell suspension was transferred to a microcentrifuge tube and sonicated on ice using a Branson 450 Digital Sonifier (20% amplitude, 20 sec total, 2.5 sec on, 2.5 sec off).

Crude lysates were centrifuged at 15,000 x g for 10 min at 4°C and clarified lysates were transferred to fresh microcentrifuge tubes on ice. Clarified lysates were normalized for total protein to 1.9 mg/mL using Bradford reagents and a BSA standard. 26.5 µL reactions composed of lysis buffer A, nucleic acid substrates, and 3.5 µL of normalized clarified lysates were assembled in PCR strip tubes, mixed by gentle pipetting, and incubated at room temperature (∼23°C) for 60 minutes. NH_4_Cl solutions at the indicated concentration were dissolved in lysis buffer A and substituted for nucleic acid substrates as positive ammonium controls.

**Ammonium Detection:** The evolution of NH_4_ by deamination of the nucleic acid substrates was observed using a phenol-hypochlorite reaction to produce indophenol in a clear 96-well microtiter plate and modified from Dong et al. 2015 [102]. The work of Ngo et al. [103] was considered when designing the lysis buffer so as not to interfere with the phenol-hypochlorite reaction. 50 µL of Reagent A (composition below) was added to each well followed by 20 µL of the completed in vitro deamination reaction described above. The phenol-hypochlorite reaction was initiated by the addition and gentle mixing of 50 µL Reagent B (composition below) to the wells. The reaction was incubated at 35°C for 30 min and the ABS_630_ was measured using a plate reader.

*Reagent A* = 1:1 (v/v), 6% (w/v) sodium hydroxide (Sigma) in water: 1.5% (v/v) sodium hypochlorite solution (Sigma, reagent grade) in water.

*Reagent B* = 1:1:0.04 (v/v/v), water: 0.5% (w/v) sodium nitroprusside (Sigma) in water: phenol solution (Sigma, P4557)

### Western Blot

Strains containing DcdV- and variant-C-terminal 6x-histidine fusions were grown, induced, and harvested as described previously above (See In-vitro Nucleic Acid Deamination Assay: Cell Lysate Prep), except for the His-tag fusion (pGBS98) which are induced for only 2 h with 100 μM IPTG and not subjected to sonication. The cell pellets were resuspended in 2 mL of chilled 1X PBS and subsequently normalized to OD of 1.0. 1 mL aliquots were collected by centrifugation at 15k x g for 1 min. Cell pellets were subsequently resuspended in 90 µL of lysis buffer A and 30 µL of 4x Laemmli buffer, denatured for 10 minutes at 65°C, and centrifuged at 15k x g for 10 minutes. 5 µL of samples were loaded into a precast 4-20% sodium dodecyl sulphate-polyacrylamide gel electrophoresis (SDS-PAGE) gels (Mini-PROTEAN TGX Precast Protein Gels, Bio-Rad) alongside size standards (Precision Protein Plus, Bio-Rad). Gels were run at room temperature for 90 min at 100 V in 1x Tris/glycine/SDS running buffer. Proteins were transferred to nitrocellulose membranes (Optitran). The membranes were blocked using 5% skim milk and incubated with 1:5000 THE^TM^ His Tag Antibody, mAb, Mouse (GenScript) followed by 1:4000 Goat Anti-Mouse IgG Antibody (H&L) [HRP], pAb (GenScript), treated with Pierce™ ECL Western Blotting Substrate, and imaged using an Amersham^TM^ Imager 600.

### UPLC-MS/MS Quantification of In Vitro and In Vivo Deoxynucleotides

Deoxynucleotide concentrations were determined as previously described [104] with minor modifications. For measuring in vivo intracellular deoxynucleotide concentrations, overnight cultures were subcultured 1:1000 and grown to OD_600_ of ∼1.0. Plasmid expression was induced by the addition of 1 mM IPTG for 1 h, and 1 mL of cultures were collected by centrifugation at 15,000 x g for 1 min. Cell pellets were resuspended in 200 µL of chilled extraction buffer [acetonitrile, methanol, ultra-pure water, formic acid (2:2:1:0.02, v/v/v/v)]. To normalize in vivo nucleotide samples, an additional cell pellet was collected from 1 mL of culture by centrifugation at 15,000 x g for 1 min, resuspended in 200 μL lysis buffer B (20 mM Tris·HCl, 1% SDS, pH 6.8), and denatured for 10 minutes at 60°C. Denatured lysates were centrifuged at 15,000 x g for 1 min to pellet cellular debris, and the supernatant was used to quantify the total protein concentration in the sample using the DC protein assay (Bio-Rad) a BSA standard curve [34]. The concentrations of deoxynucleotides detected by UPLC-MS/MS were then normalized to total protein in each sample.

For the quantification of deoxynucleotides in vitro *E. coli* BL21(DE3) clarified lysates were prepared as described for the deamination experiment above and normalized to 20 mg/mL of total protein and 200 µL of normalized clarified lysates were assembled in PCR strip tubes.

To measure abundance of dUMP and dUTP prior to the addition of 1 µM dCTP, 20 µL of normalized clarified lysates were added to 200 µL of chilled extraction buffer. 20 µL of 10 µM dCTP was then added to the remaining clarified lysates and 20 µL lysates aliquots were removed 1, 5, 10, and 30 minutes after the addition of dCTP and mixed in 200 µL chilled extraction buffer.

All samples resuspended in extraction buffer, in vivo and in vitro, were immediately incubated at -20°C for 30 minutes after collection and centrifuged at 15,000 x g for 1 min. The supernatant was transferred to a new tube, dried overnight in a speed vacuum, and finally resuspended in 100 μL ultra-pure water. Experimental samples and deoxynucleotides standards [1.9, 3.9, 7.8, 15.6, 31.3, 62.5, and 125 nM of dATP (Invitrogen), dGTP (Invitrogen), dTTP, (Invitrogen), dCTP (Invitrogen), dCMP (Sigma), dUTP (Sigma), and dUMP (Sigma)] were analyzed by UPLC-MS/MS using an Acquity Ultra Performance LC system (Waters) coupled with a Xevo TQ-S mass spectrometer (Waters) with an ESI source in negative ion mode. The MS parameters were as follows: capillary voltage, 1.0 kV; source temperature, 150°C; desolvation temperature, 400°C; cone gas, 120 L/hr. Five microliter of each sample was separated in reverse phase using Acquity UPLC Premier BEH C18, 2.1 x 100 mm, 1.7 µm particle size, VanGuard FIT at a flow rate of 0.3 mL/min with the following gradient of solvent A (8mM DMHA (N,N-dimethylhexylamine) + 2.8 mM acetic acid in water, pH∼9) to solvent B (methanol): *t* = 0 min; A-100%:B-0%, *t* = 10 min; A-60%:B-40%, *t* =10.5; A-100%:B-0%, *t* = 15 min; A-100%:B-0% (end of gradient). The conditions of the MRM transitions were as follows [cone voltage (V), collision energy (eV)]: dATP, 490 > 159 (34, 34); dCTP, 466 > 159 (34, 34); dGTP, 506 > 159 (15, 46); dTTP, 481 > 159 (25, 34); dUTP, 467 > 159 (25, 34); dCMP, 306 > 97 (43, 22); dUMP, 306 > 111 (22, 22).

### Phage Infection and Plaque Assays

*V. cholerae* phages ICP1 and ICP3 were provided by Wai-Leung Ng at Tuft University School of Medicine. ICP1 was propagated on *V. cholerae* E7946, while ICP3 were propagated on *V. cholerae* C6706str2 in LB, and their titer was determined using the small drop plaque assay method, as previously described [10]. Briefly, 1 ml of overnight cultures were mixed with 9 ml of MMB agar (LB + 0.1 mM MnCl2 + 5 mM MgCl2 + 5 mM CaCl2 + 0.5% agar), tenfold serial dilutions of phages in MMB were dropped on top of them, and incubated overnight at 35°C. The viral titer is expressed as plaque forming units per mL (pfu/mL). 4 mL of *V. cholerae* overnight cultures were diluted 1:1000 in MMB medium. 145 µL of the diluted cultures, in three sets of biological replicates, were transferred and incubated at 35°C in a 96-well microplate (Costar®). Once the OD_600_ reached ∼0.1, 5 µL of phages with a final MOI of 0.1 were added to each biological replicate. Cultures were infected at room temperature (∼23°C) for 12 h in a SpectraMax M5 Plate Reader with continuous shaking and OD_600_ measurements taken every 2.5 min.

*Shigella flexneri* strain PE577 [54] cells transformed with the pVector (pMMB67eh) and each of the associated pDcdV plasmids were grown in LB medium and incubated with aeration at 37° C overnight. The following day, 20 µL of each of the overnight cultures were used to inoculate fresh medium in a 96-well microtiter plate with a final volume of 200 µL/well.

Depending on the experimental condition, wells were supplemented with and without IPTG (100 µM final concentration) and/or phage Sf6 [55] at an MOI of 0.1 phage per cell. Initial cell densities of the overnight cultures were experimentally determined by plating and found to be within a factor of two of one another. For all experiments, three biological replicates were tested. Additionally, the plates were set up with each unique condition having three technical replicates. Plate reader assays were conducted using a Molecular Devices FilterMax F5 plate reader, as previously described [105]. Briefly, the plates were incubated at 37°C for 6 h. Every five minutes, the plate was mixed and aerated by orbital shaking before an absorbance (595 nm) reading was taken. After the kinetic assay was complete an aliquot from each of the replicates was removed and used to determine the endpoint titer via plaque assay.

### Statistical Analysis

As specified in the figure legends, all of the statistical analyses for the violin plots were performed with R statistical computing software [90], while other data were analyzed in GraphPad Prism Software. Statistically significances denote as the following: a single asterisk (*) indicates p < 0.05; double asterisks (**) indicate p < 0.01; triple asterisks (***) indicate p < 0.001; and quadruple asterisks (****) indicate p < 0.0001. Means ± SEM and specific n values are reported in each figure legend.

## Graphical Abstract

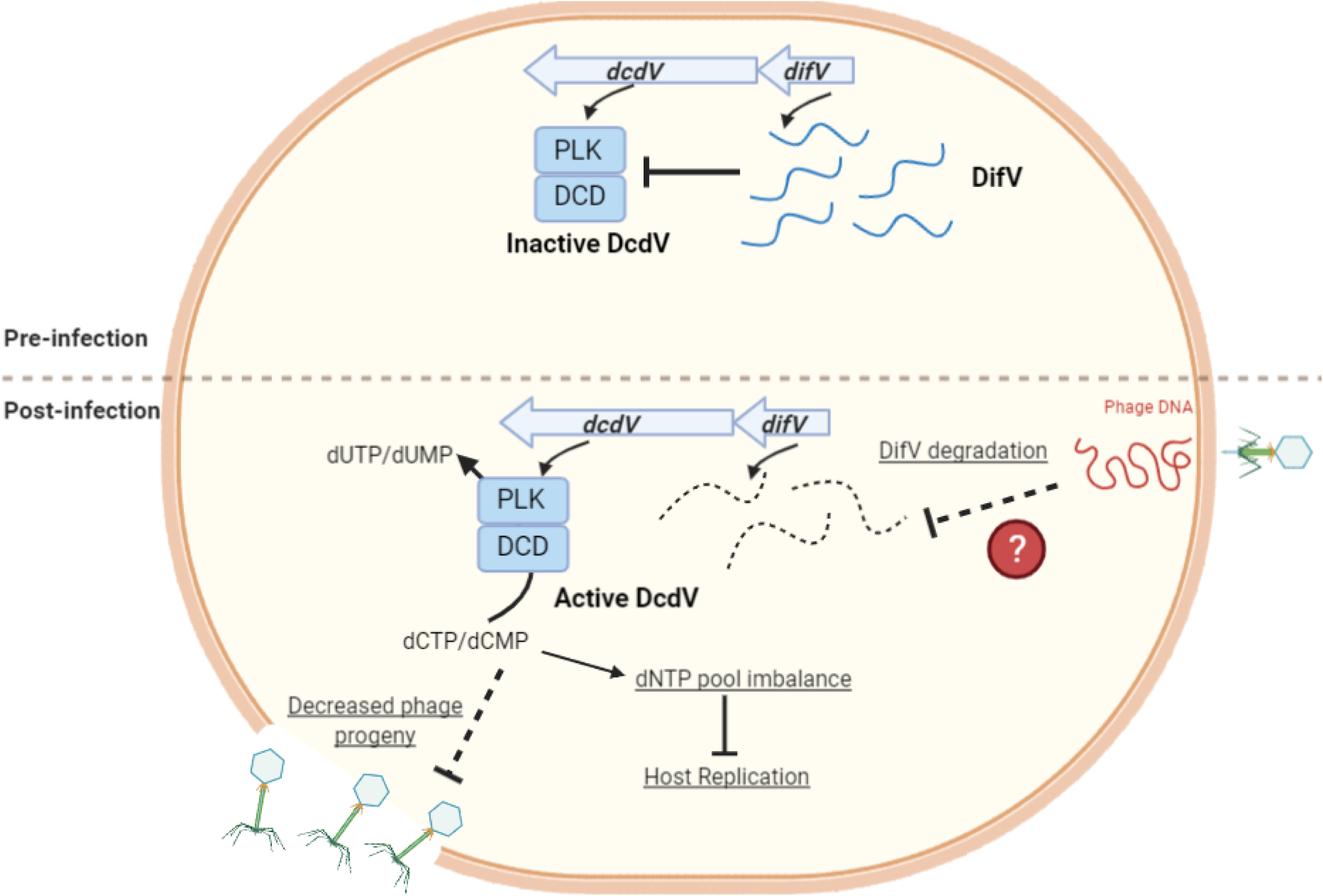

## SUPPLEMENTAL MATERIAL

**Supplemental Figure 1.**
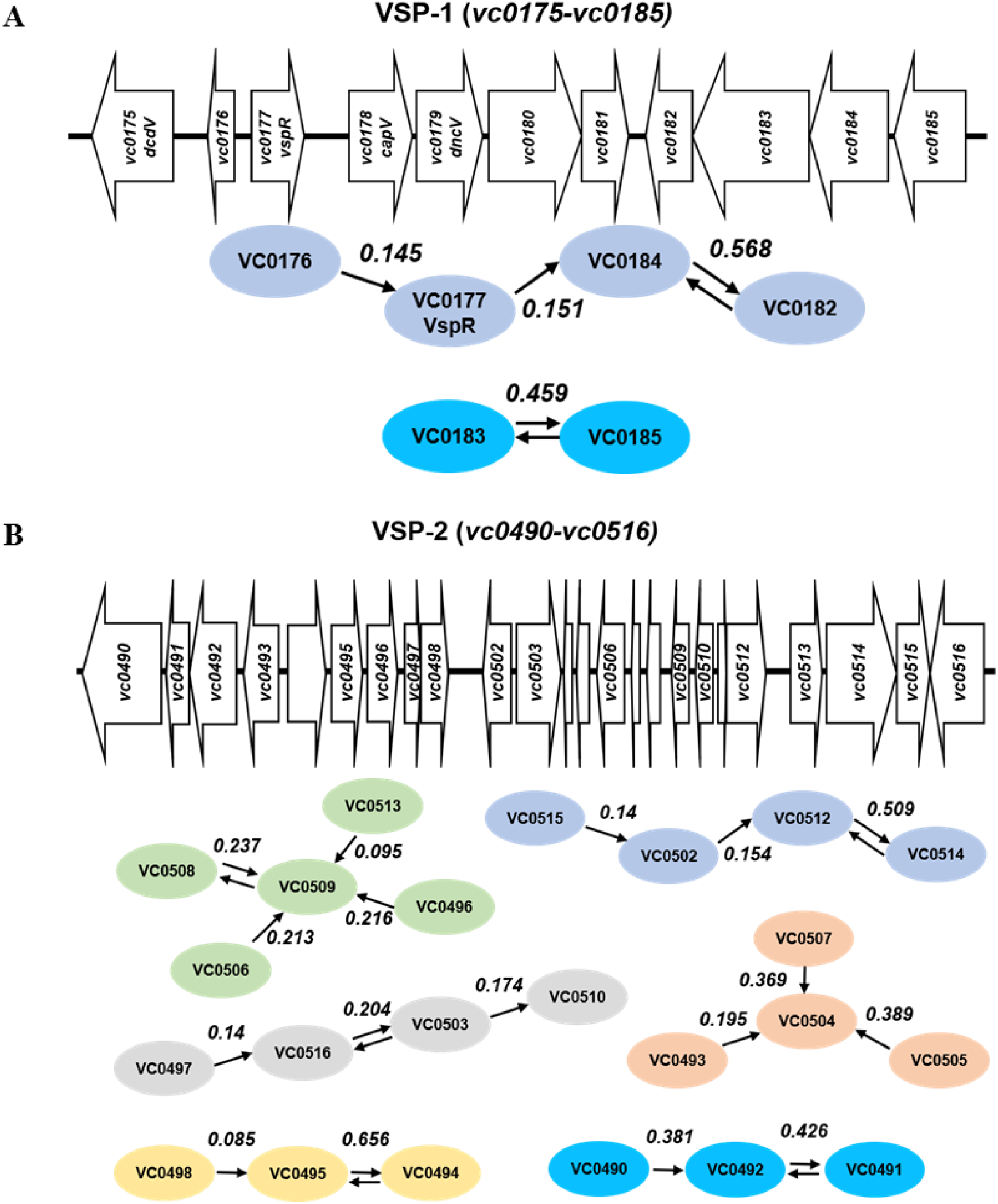
VSP-1 and VSP-2 schematic and predicted maximum related subnetworks (MRS). (**A**) Cartoon schematic and gene network predictions, other than DcdV and CBASS (see Figs. 1B and 1), of VSP-1 from El Tor *V. cholerae* N16961 (not to scale). (**B**) Cartoon schematic and gene network predictions of VSP-2 from El Tor *V. cholerae* N16961 (not to scale). Arrows indicate the highest partial correlation ***W_ij_*** of each individual VSP gene to another (represented by ovals). Two arrows pointing in opposing directions indicates the two genes each have their highest correlation to each other.

**Supplemental Figure 2.**
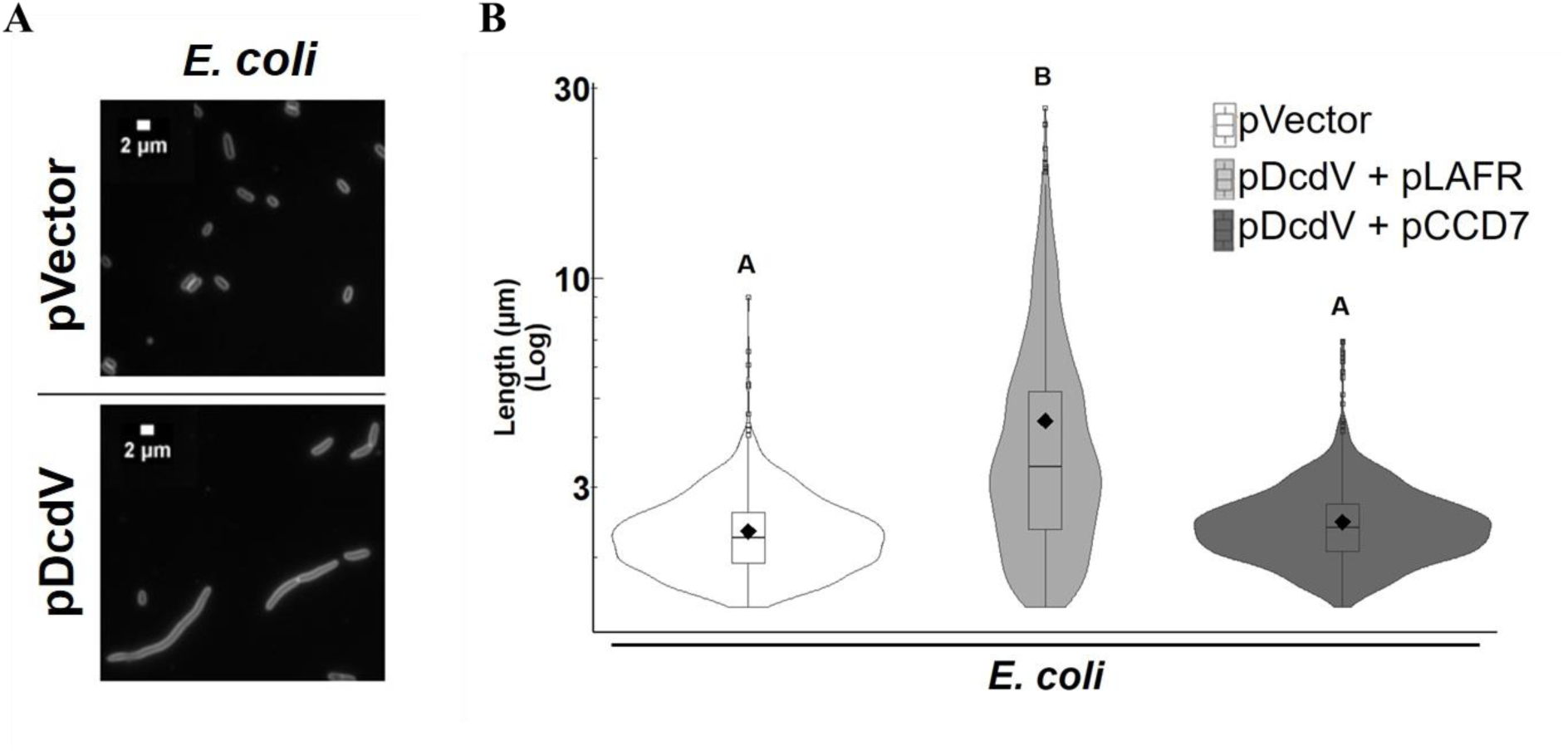
Ectopic expression of *dcdV* leads to cell filamentation in *E. coli* that is alleviated by provision of a single copy cosmid containing VSP-1. (**A**) Representative images of *E. coli* cultures maintaining an empty vector plasmid (pVector) or P_tac_-inducible *dcdV* plasmid (pDcdV) grown in the presence of 100 µM IPTG for 8 h. Cells were stained with FM4-64 prior to imaging. Scale represents 2 µm. (**B**) Distribution of cell lengths measured from three biological replicates of *E. coli* cultures carrying an empty vector (Vector) or P_tac_-inducible *dcdV* plasmid (pDcdV) in addition to either an empty vector single copy cosmid control (pLAFR) or pLAFR containing VSP-1 (pCCD7) grown in the presence of 100 µM IPTG for 8 h. Distributions represent ∼1000 to 2000 cells measured per strain. Different letters indicate significant differences at *p* < 0.05, according to Tukey’s post-hoc test.

**Supplemental Figure 3.**
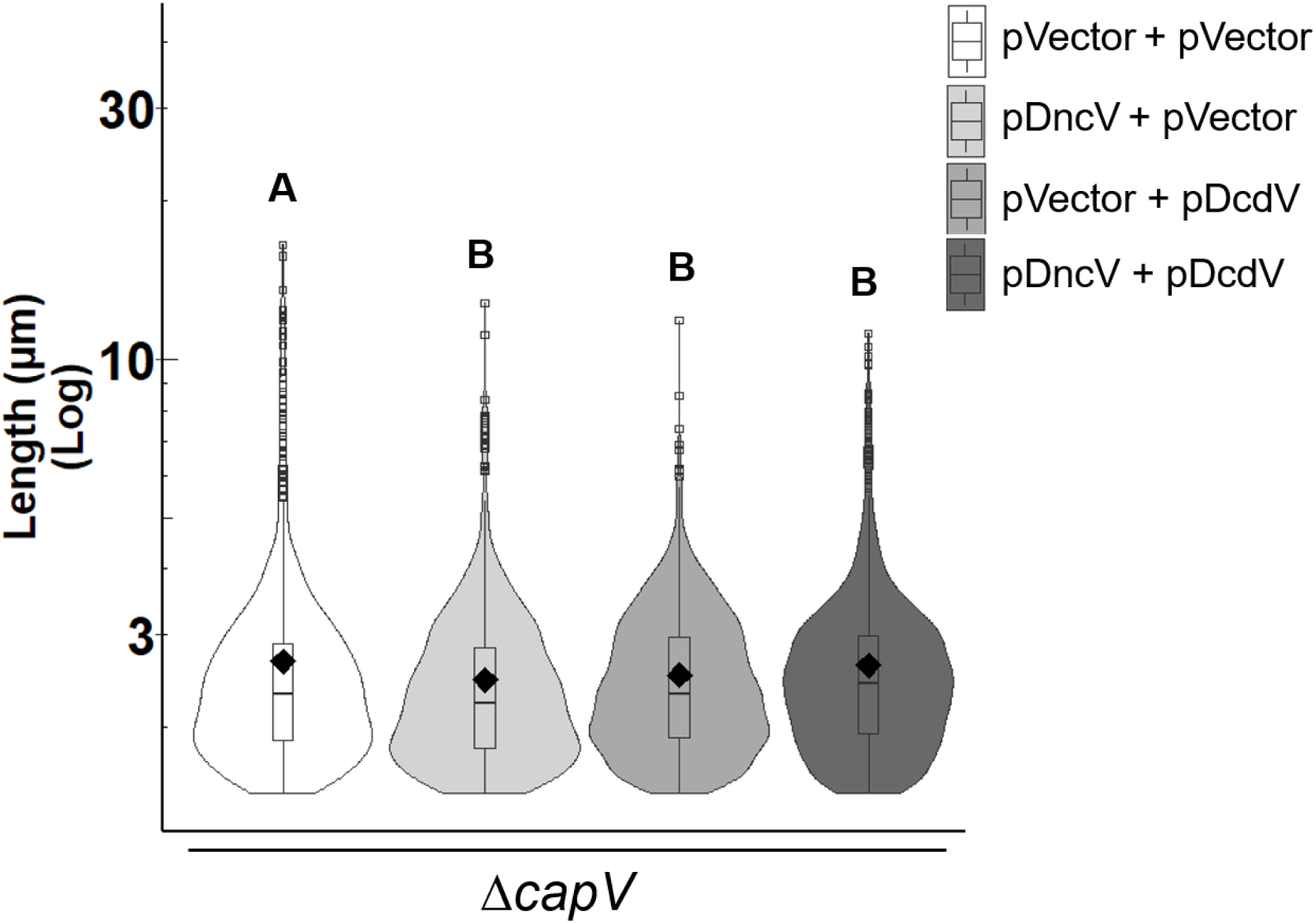
Ectopic expression of DncV and DcdV does not lead to filamentation in the Δ*capV* mutant of *V. cholerae.* Distribution of cell lengths measured from three biological replicates of *ΔcapV* mutant cultures maintaining the indicated plasmids grown in the presence of 100 µM IPTG for 8 h. Distributions represent ∼1200-1700 cells measured per strain. Different letters indicate significant differences at *p* < 0.05, according to Tukey’s post-hoc test.

**Supplemental Figure 4.**
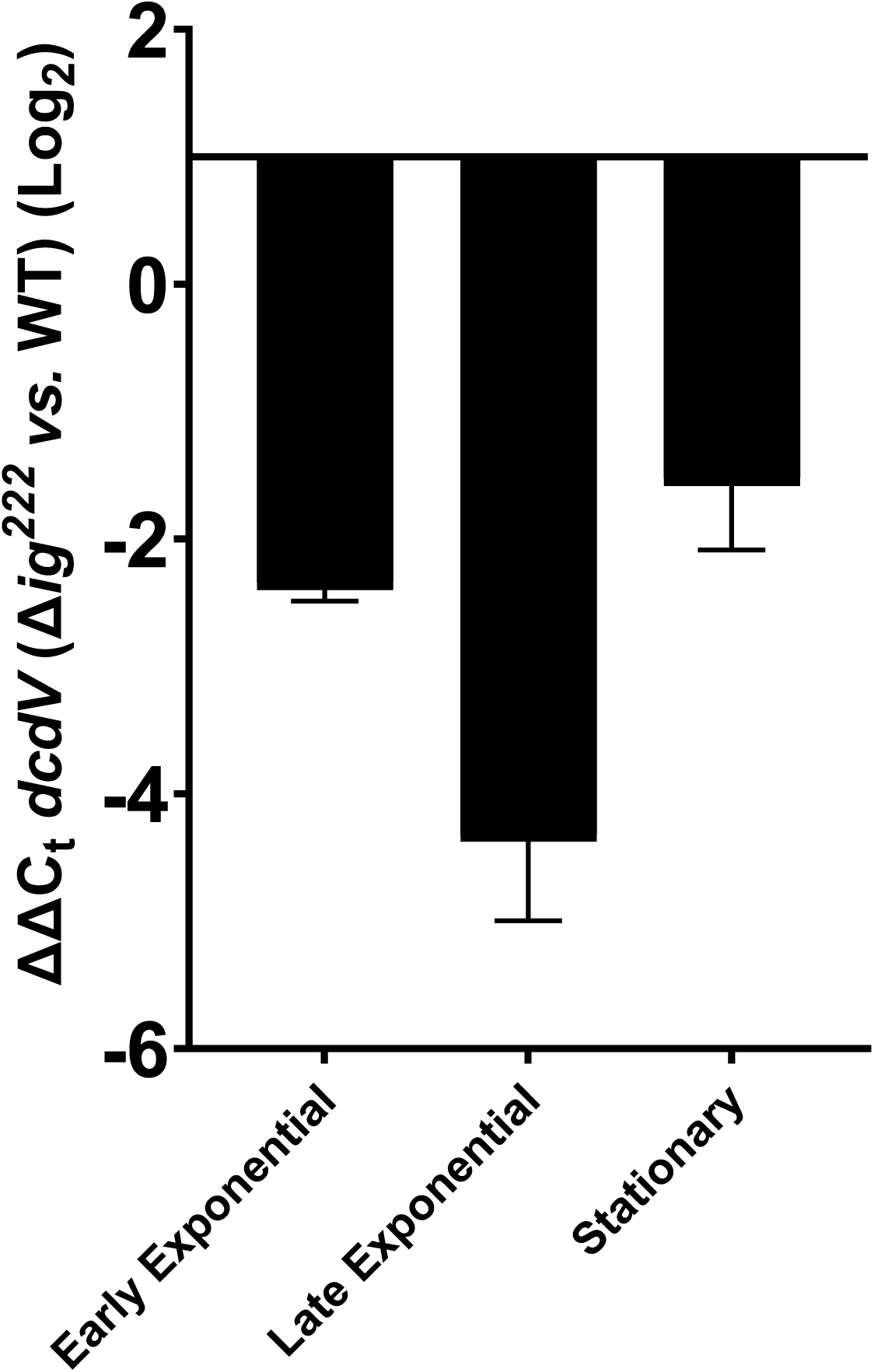
Δig222 has decreased *dcdV* expression relative to WT *V. cholerae*. Relative difference in *dcdV* expression between Δ*ig^222^* and WT *V. cholerae* at three different growth phases using qRT-PCR and an endogenous *gyrA* control. Data represent the mean ± SEM of three biological replicates.

**Supplemental Figure 5.**
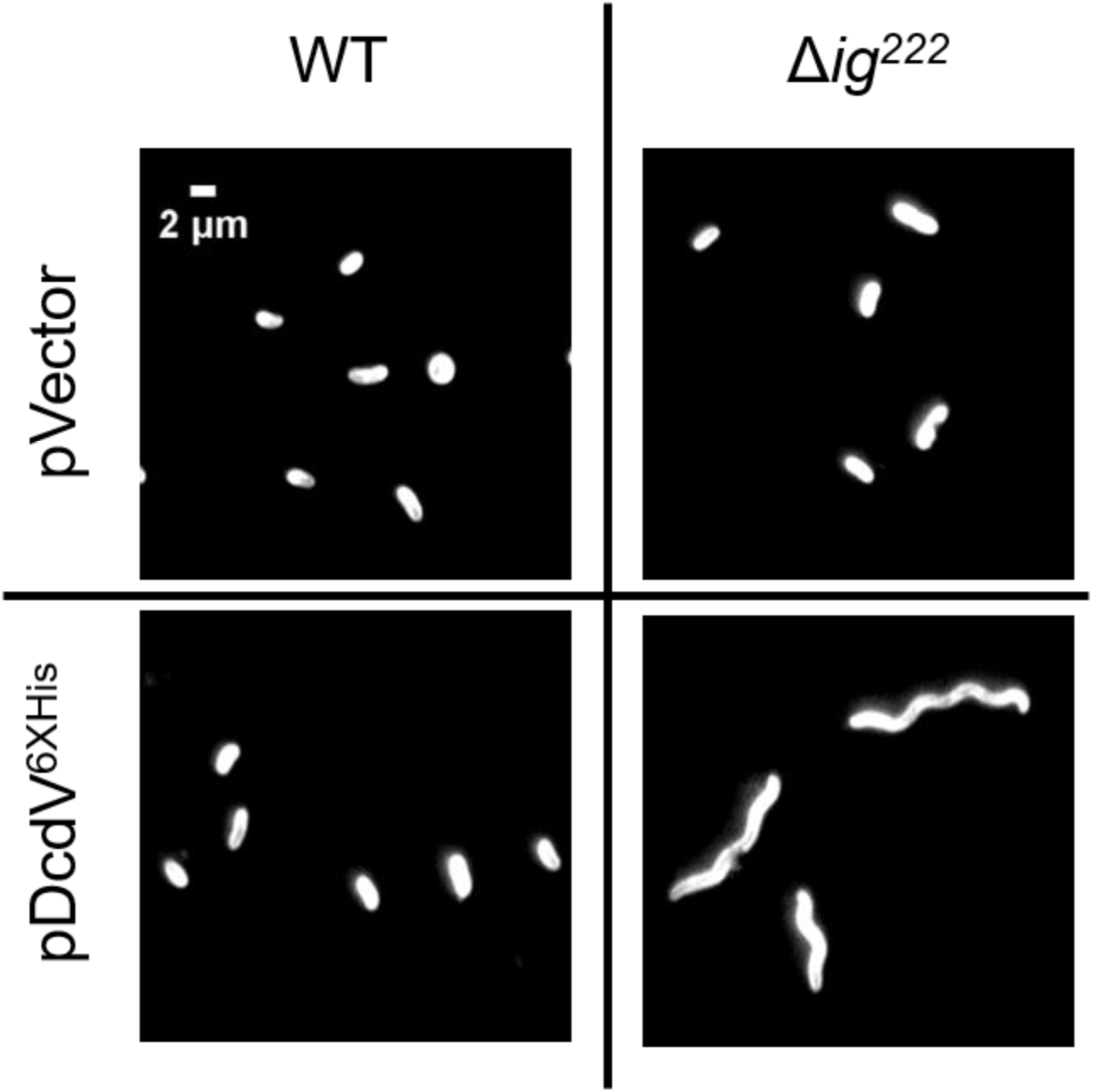
DcdV C-terminal 6x Histidine fusion maintains the same activity as the WT DcdV enzyme. Representative images of *V. cholerae* WT and Δ*ig^222^* cultures maintaining an empty vector plasmid (pVector) or P_tac_-inducible *dcdV-6xHIS* plasmid (pDcdV^6xHis^) grown in the presence of 100 µM IPTG for 2 h. Cells were stained with FM4-64 prior to imaging and performed in biological triplicate.

**Supplemental Figure 6.**
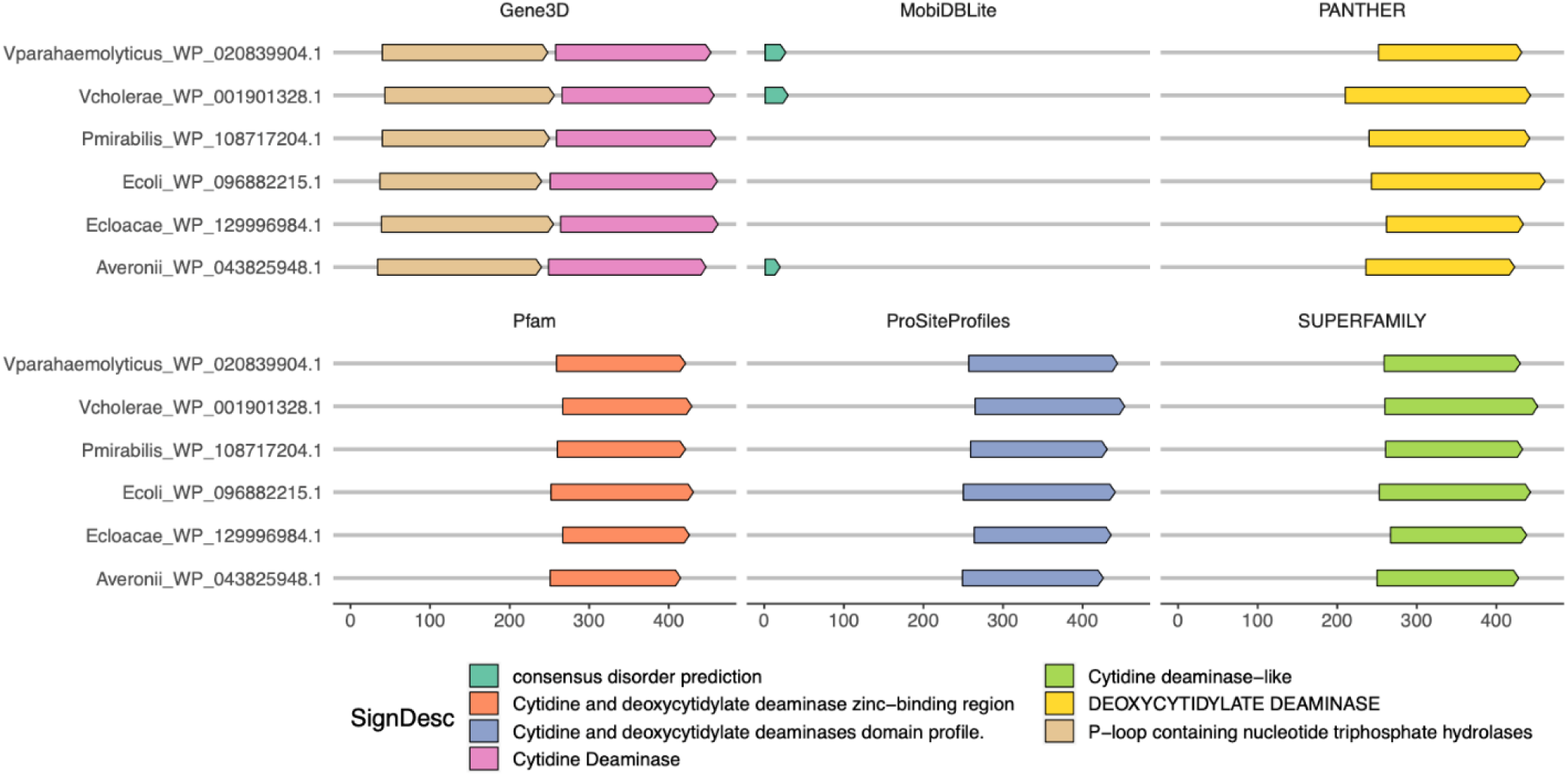
Domain architectures of the six DcdV query proteins. Domain architecture and secondary structure predictions for the six proteobacterial starting points of interest (query proteins) using InterProScan [[40]; see Methods]. Results from six main analyses are shown here for the query proteins: Gene3D (including CATH structure database), Pfam, ProSiteProfiles, PANTHER, and SUPERFAMILY protein domain profile databases, and MobiDBLite for disorder prediction. No transmembrane regions (using TMHMM) or membrane/extracellular localization were predicted for any of the proteins (using Phobius); hence not shown.

**Supplemental Figure 7.**
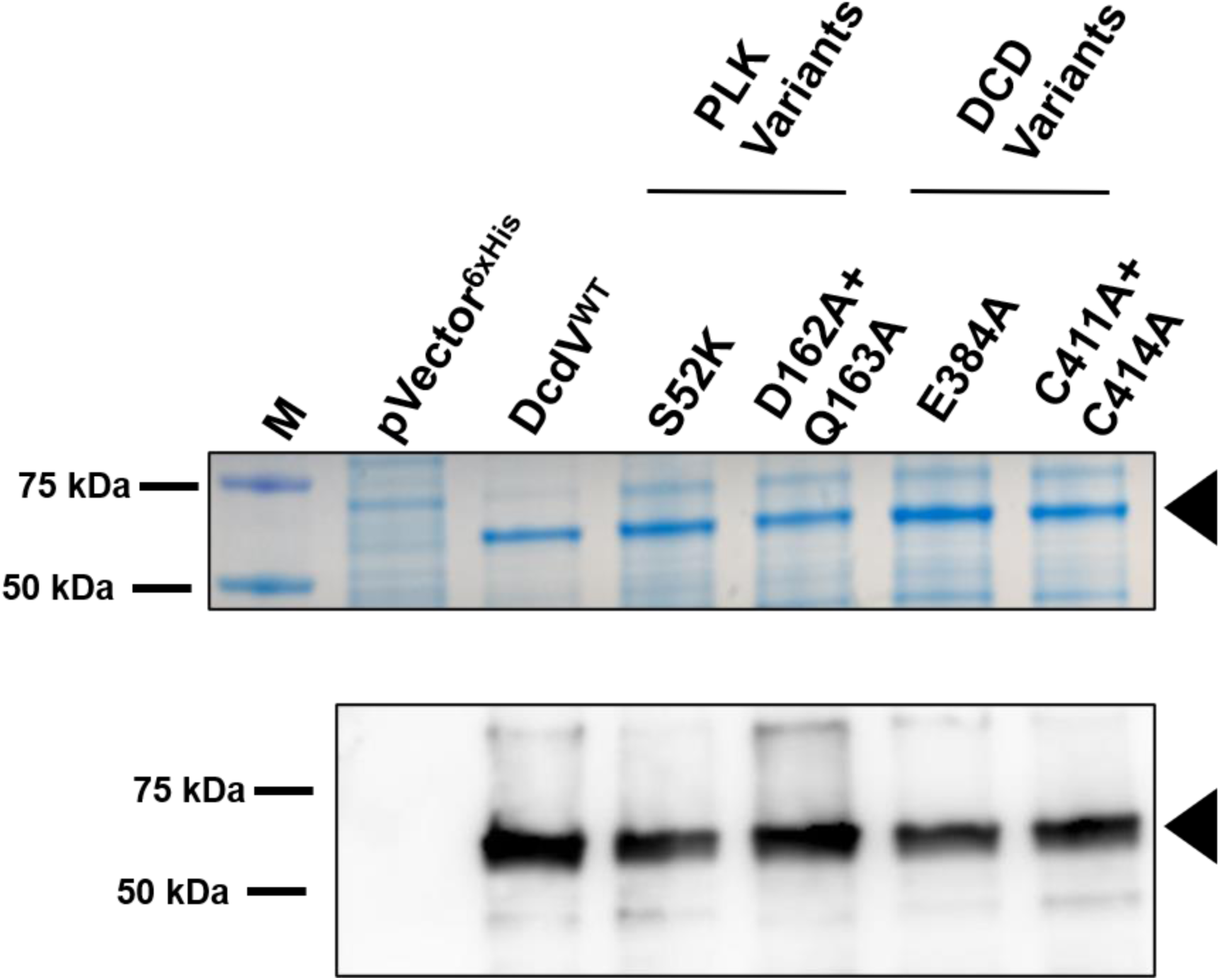
Cellular abundance of C-terminal 6x histidine tagged DcdV variant fusions analyzed by Coomassie stain and Western blot. Representative Coomassie stained gel (top) and anti-6x His antibody Western blot (bottom) of whole cell lysates from *E. coli* BL21(DE3) cells maintaining an empty vector (pVector^6xHis^), inducible C-terminal 6x histidine tagged *dcdV* (WT) or *dcdV* variants (S52K, D162A + Q163A, E384A, and C411A + C414A) grown in the presence of 1 mM IPTG for 3 h. Sample inputs were normalized by culture OD_600_ and resolved by SDS-PAGE. Three biological replicates of each strain were analyzed with similar results. Black triangles correspond to the predicted molecular weight of the DcdV tagged fusions (60.6 kDa). M = molecular weight marker.

**Supplemental Figure 8.**
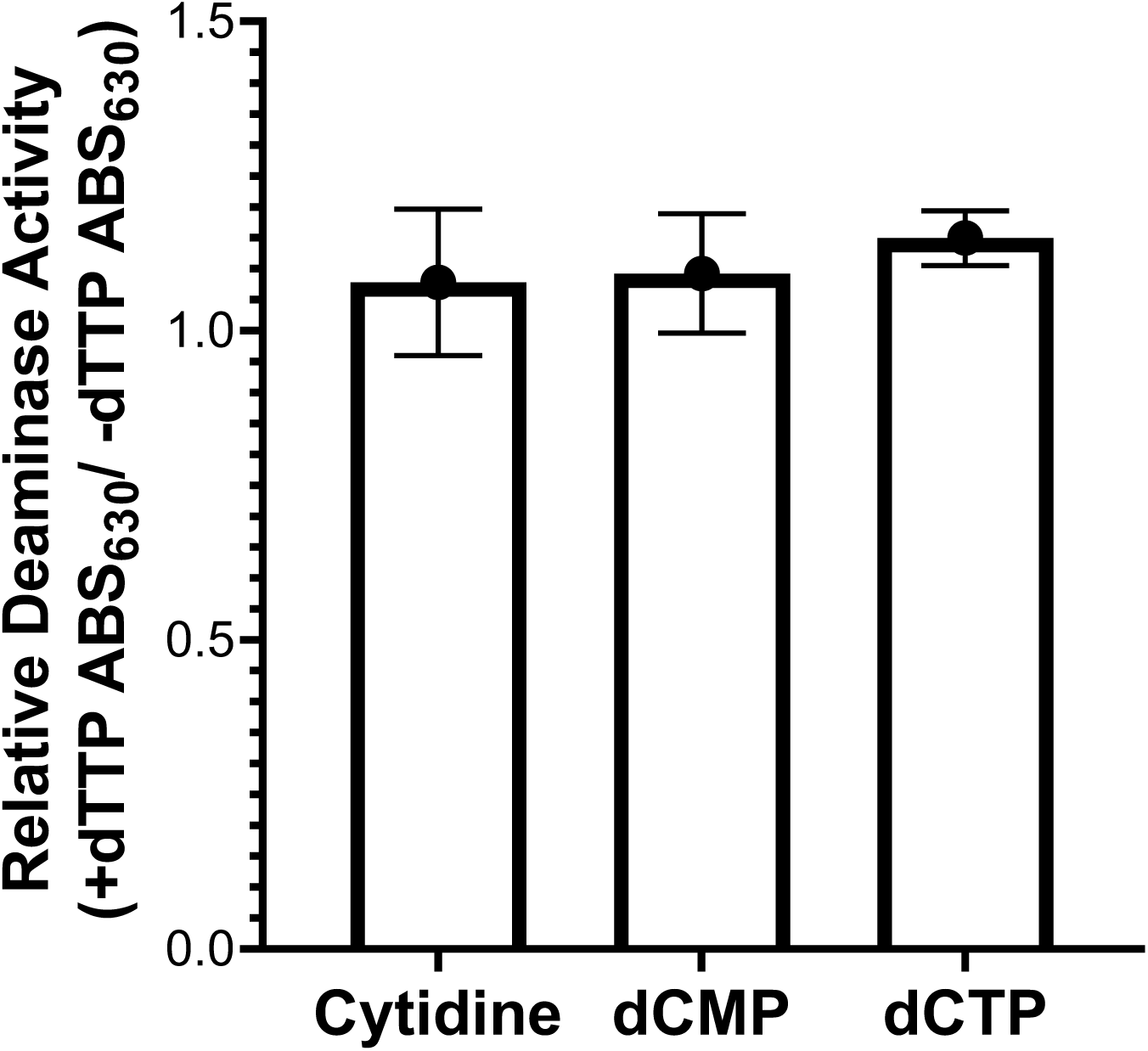
Addition of exogenous dTTP does not inhibit DcdV deaminase activity in *E. coli* lysates. Lysates collected from *E. coli* expressing WT DcdV incubated with or without exogenous 7.5 mM dTTP and either 75 mM cytidine, 7.5 mM dCMP, or 7.5 mM dCTP. The evolution of NH_4_^+^ resulting from substrate deamination was detected by measuring the solution ABS_630_ after a Berthelot’s reaction in microtiter plates. The relative deaminase activity was calculated by dividing the ABS_630_ of the +dTTP reaction by the no dTTP control reaction for each lysate. Data represent the mean ± SEM of three biological replicate lysates.

**Supplemental Figure 9.**
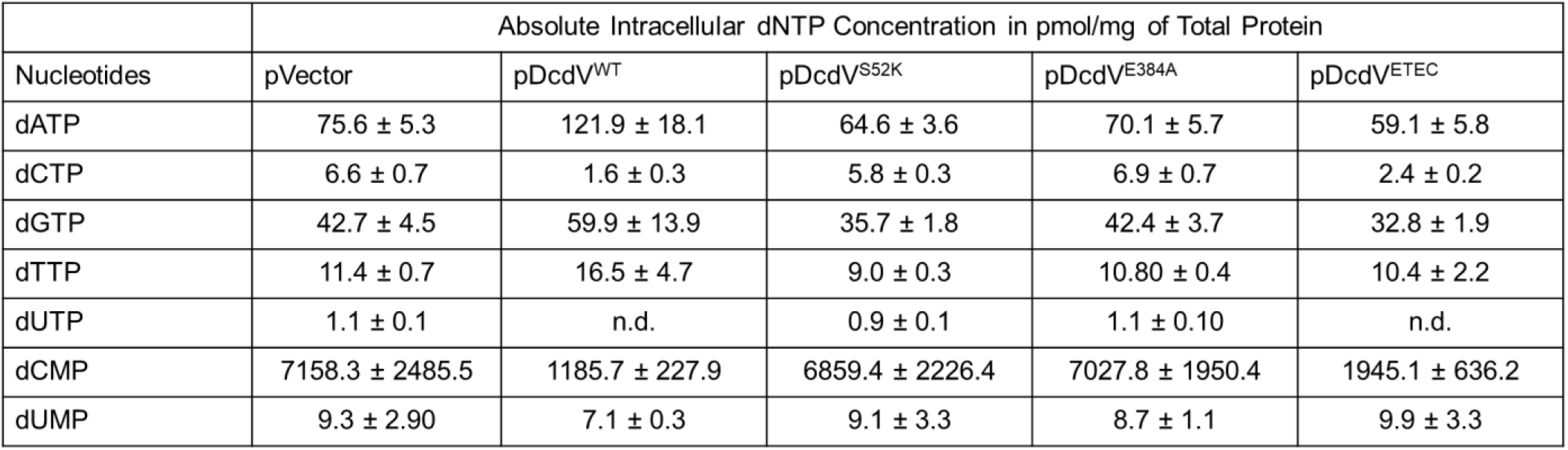
Absolute intracellular concentration of deoxynucleotides. Quantification of the indicated dNTPs in vivo, using UPLC-MS/MS, in strains expressing the empty vector and four DcdV variants, as indicated. Data represents mean ± SEM, *n*=3.

**Supplemental Figure 10.**
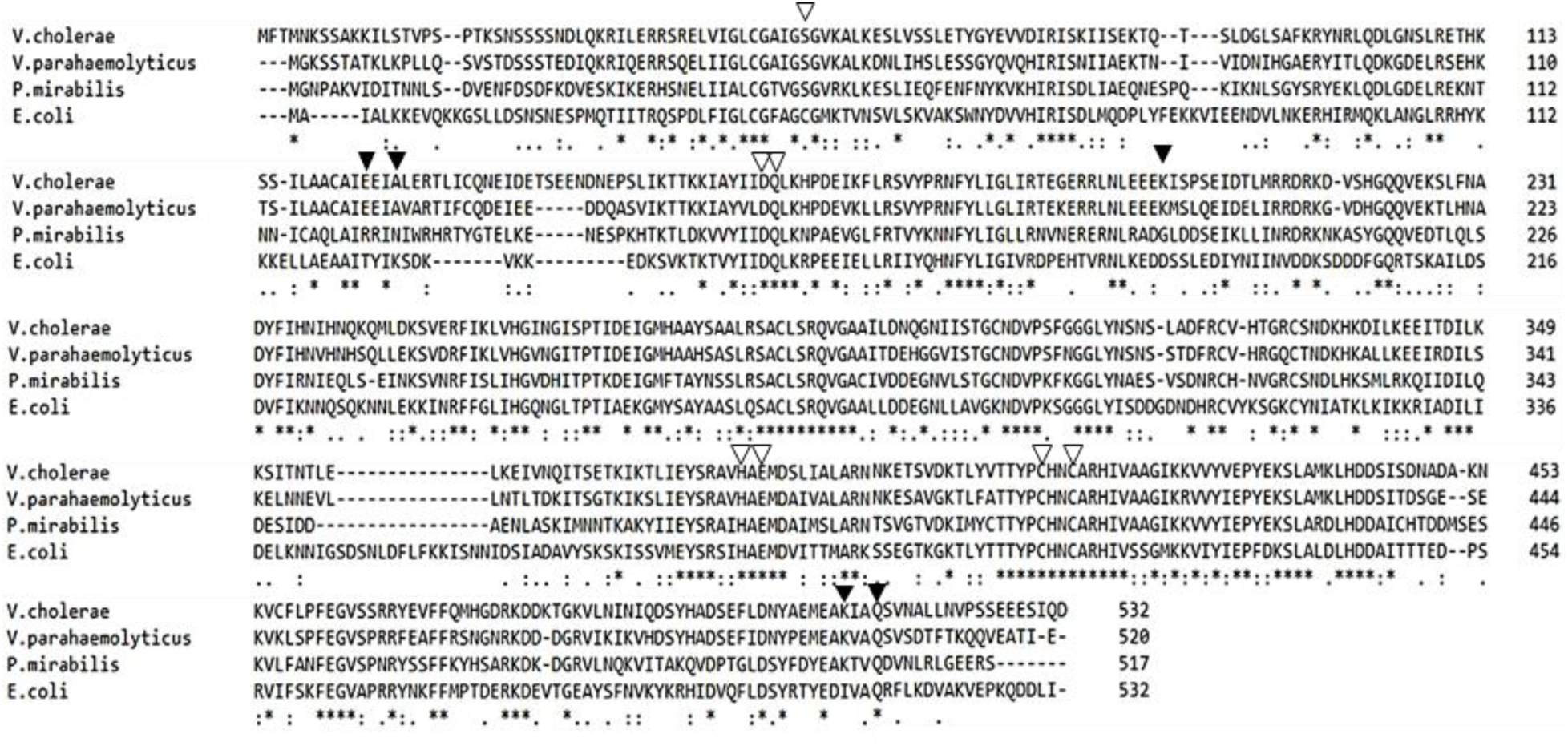
ClustalW multiple sequence alignment of DcdV homologs explored in this study. Amino acid alignment of DcdV and three homologs using webservice EMBL-EBI [74]. “**\***” indicates 100% identity, “:” indicates >75%, and “.” Indicates >50% similarity. Open triangles above the alignments indicate conserved residues of PLK and DCD domains. Closed triangles indicate amino acids where single amino acid substitutions were found to render *V. cholerae* DcdV insensitive to DifV inhibition (Figs. 6A, B, and D).

**Supplemental Figure 11.**
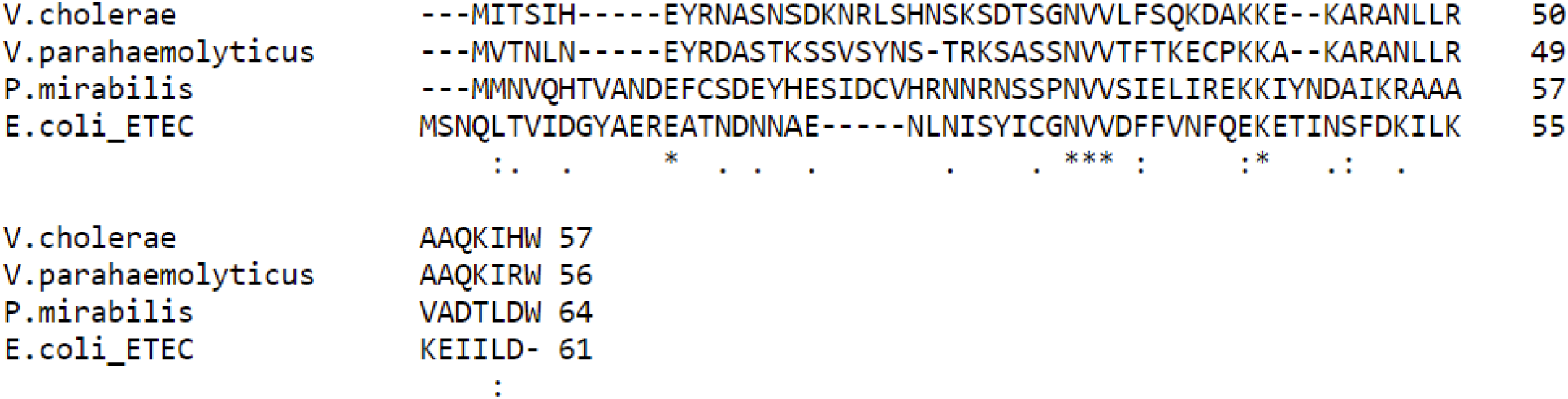
DifV (174 nt) and the three ORFs encoded upstream of *dcdV* homologs do not exhibit amino acid similarity. Amino acid alignment of the *V. cholerae* Ig^222^ translated ORF and three ORFs 5’ of the *dcdV* homologs using EMBL-EBI ClustalW [74]. “**\***” indicates 100% identity, “:” indicates >75%, and “.” Indicates >50% similarity.

**Supplemental Figure 12.**
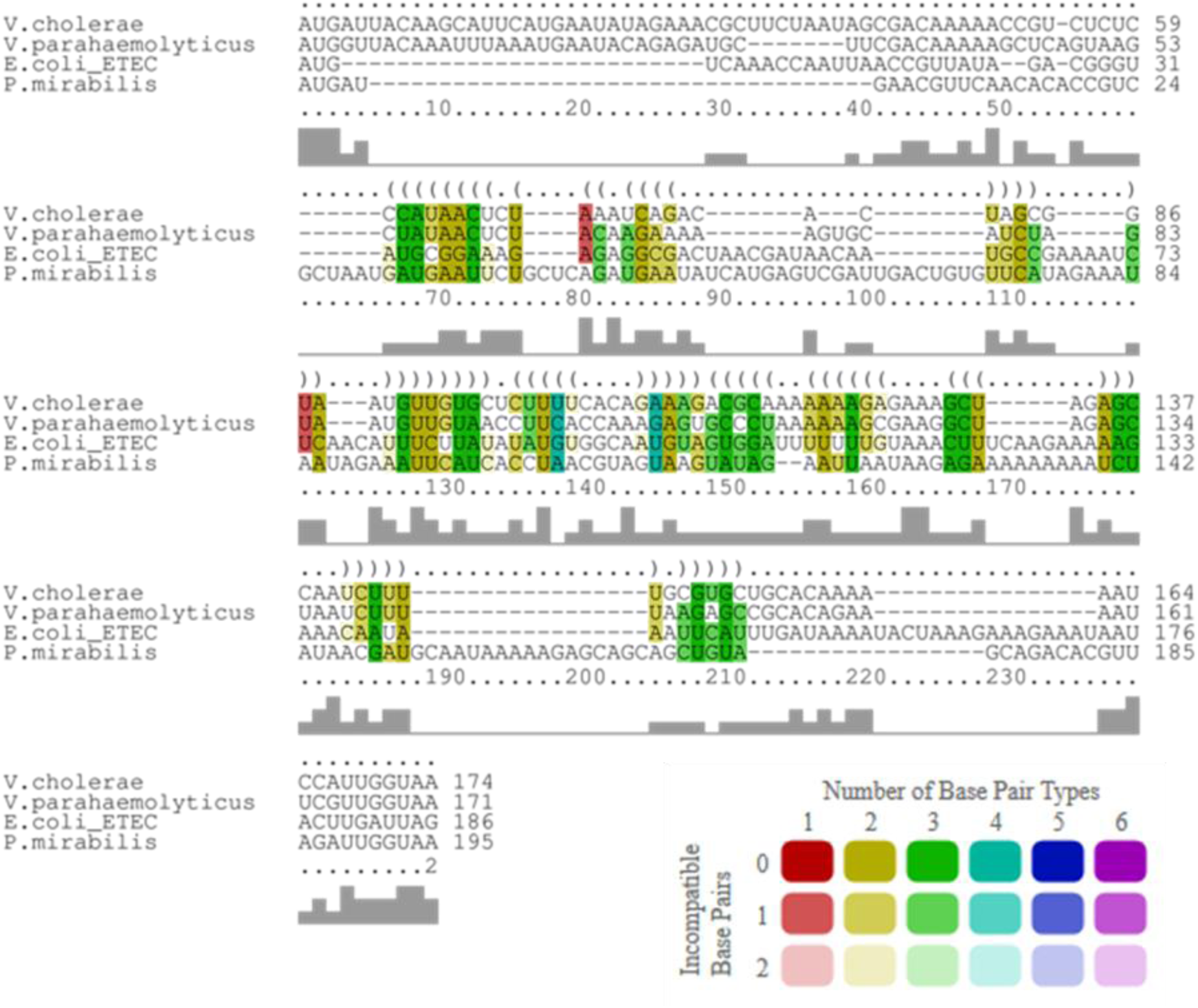
DifV (174 nt) and the three ORFs encoded upstream of *dcdV* homologs do not have exhibit similarity. Nucleotide alignment of the *V. cholerae* DifV and the ORFs 5’ of *dcdV* homologs using LocARNA [75]. Consensus identities are correlated with the height of the bars below the corresponding nucleotide (bottom). The average secondary structure is indicated in dot-bracket notation (top). Compatible base pairs are colored according to the number of different types C-G (1), G-C (2), A-U (3), U-A (4), G-U (5) or U-G (6) of compatible base pairs in the corresponding columns. The saturation decreases with the number of incompatible base pairs.

**Supplemental Figure 13.**
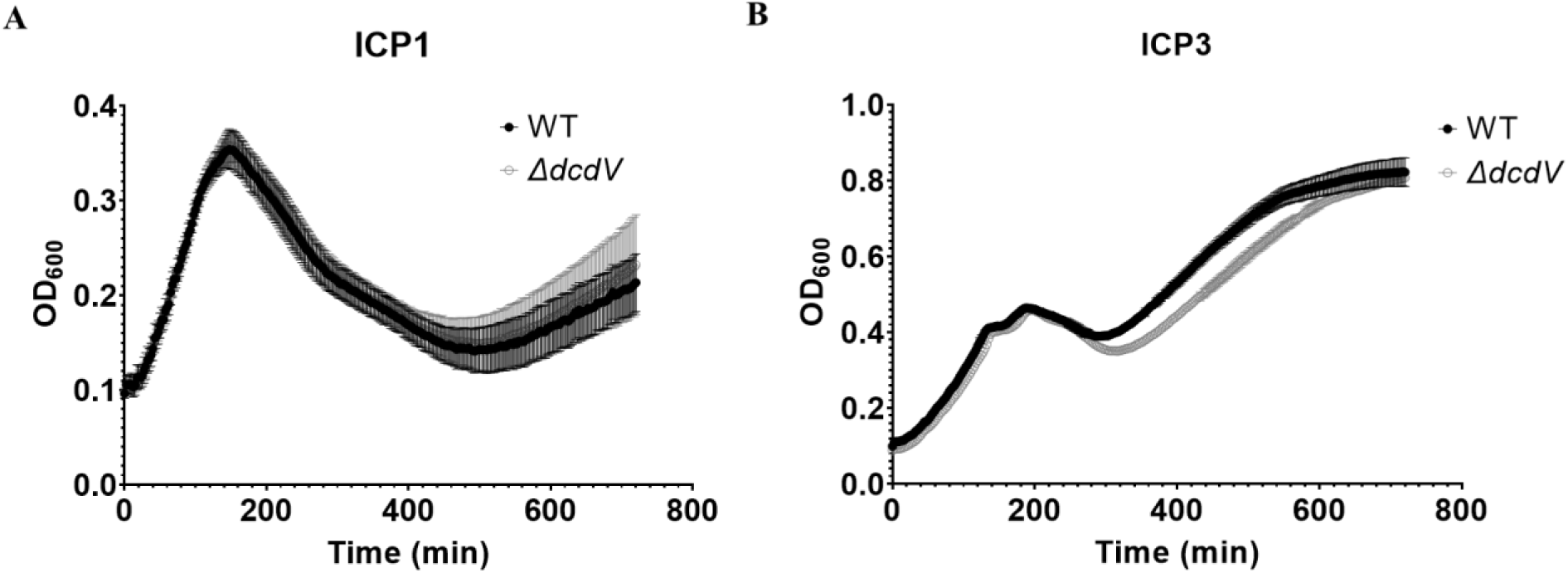
*V. cholerae* lacking *dcdV* do not exhibit enhanced susceptibility to predation by *V. cholerae* lytic phage ICP1 and ICP3. Growth curves for *V. cholerae* WT and Δ*dcdV* infected by lytic phage ICP1 (**A**) and ICP3 (**B**). Bacteria were infected at time 0 at an MOI of 0.1 in microtiter plates. Data represent the mean ± SEM, *n*=3.

**Supplementary Table 1.** Strains and phages used in this study.

| Strains | Name in this Study | Relevant Characteristics | Source or reference |
| --- | --- | --- | --- |
| ***E. coli*** |  |  |  |
| DH10b |  | *F-mcrA Δ(mrr-hsdRMS-mcrBC) Φ80lacZΔM15 ΔlacX74 recA1 endA1 araD139Δ(ara, leu)7697 galU galK λrpsL nupG* | ThermoFisher Scientific |
| BW29427 |  | *RP4-2(TetSkan1360::FRT), thrB1004, lacZ58(M15), ΔdapA1341::[erm pir<sup>+</sup>], rpsL(strR), thi-, hsdS-, pro-* | Lab Stock |
| BL21(DE3) |  | *F- ompT hsdSB(rB -mB +) gal dcm (DE3)* | Lab Stock |
| 078:H11 H10407 (ETEC) | ETEC | Wild type | [106] |
| ***V. cholerae*** |  |  |  |
| C6706str2 | WT | Wild type O1 El Tor; Sm<sup>R</sup> | [107] |
| E7946 |  | Clinical isolate obtained in Bahrain in 1978; Sm<sup>R</sup> | [108] |
| CR01 | ΔVSP-1 | O1 El Tor ΔVSP-1 | This study |
| CR02 | ΔVSP-2 | O1 El Tor ΔVSP-2 | This study |
| CR03 | ΔVSP-1/2 | O1 El Tor ΔVSP-1/2 | This study |
| BYH206 | Δ*ig*<sup>222</sup> | O1 El Tor Δ*ig*<sup>222</sup> between *vc0175-vc0176* position in N16961 chromosome I [177,230-177,008] | This study |
| BYH207 | Δ*vc0176* | O1 El Tor Δ*vc0176* | This study |
| BYH255 | Δ*vc0175-176* | O1 El Tor Δ*vc0175-176* | This study |
| BYH256 | Δ*vc0177-181* | O1 El Tor Δ*vc0177-181* | This study |
| BYH257 | Δ*vc0182-185* | O1 El Tor Δ*vc0182-185* | This study |
| GS05 | Δ*vc0175* | O1 El Tor Δ*vc0175* (*dcdV*) | This study |
| WLN5105 | Δ*capV* | O1 El Tor Δ*capV* | [11] |
| ***V. parahaemolyticus*** |  |  |  |
| O1:Kuk str. FDA_R31 | VP | Wild type | [109] |
| ***P. mirabilis*** |  |  |  |
| AR379 | PM | Wild type | [110] |
| ***Shigella flexneri*** |  |  |  |
| PE577 | Sf | Wild type | [54] |
| **Phages** |  |  |  |
| ICP1 | ICP1 | Wild type | [49] |
| ICP3 | ICP3 | Wild type | [49] |
| Sf6 | Sf6 | Wild type | [111] |

**Supplementary Table 2.**
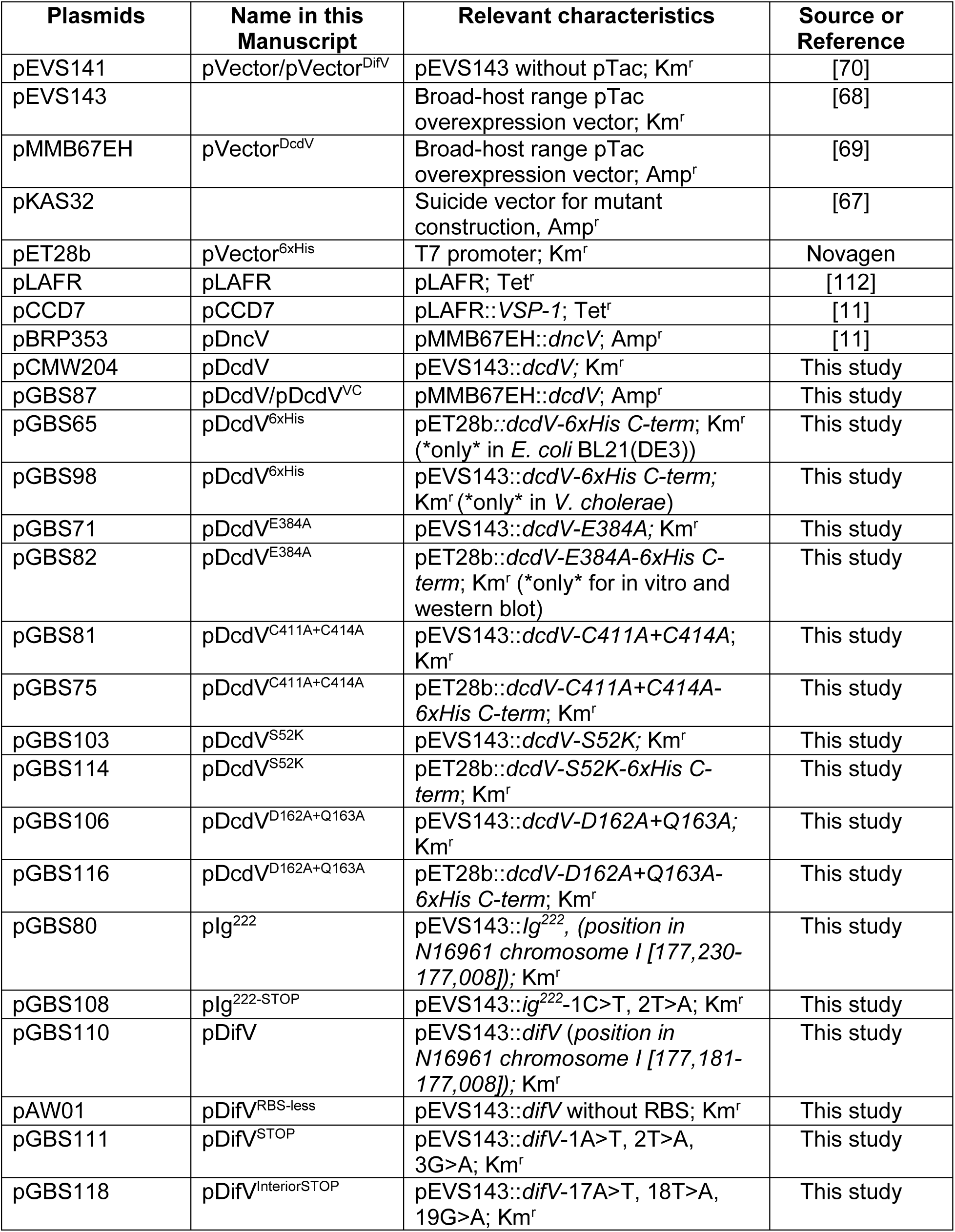
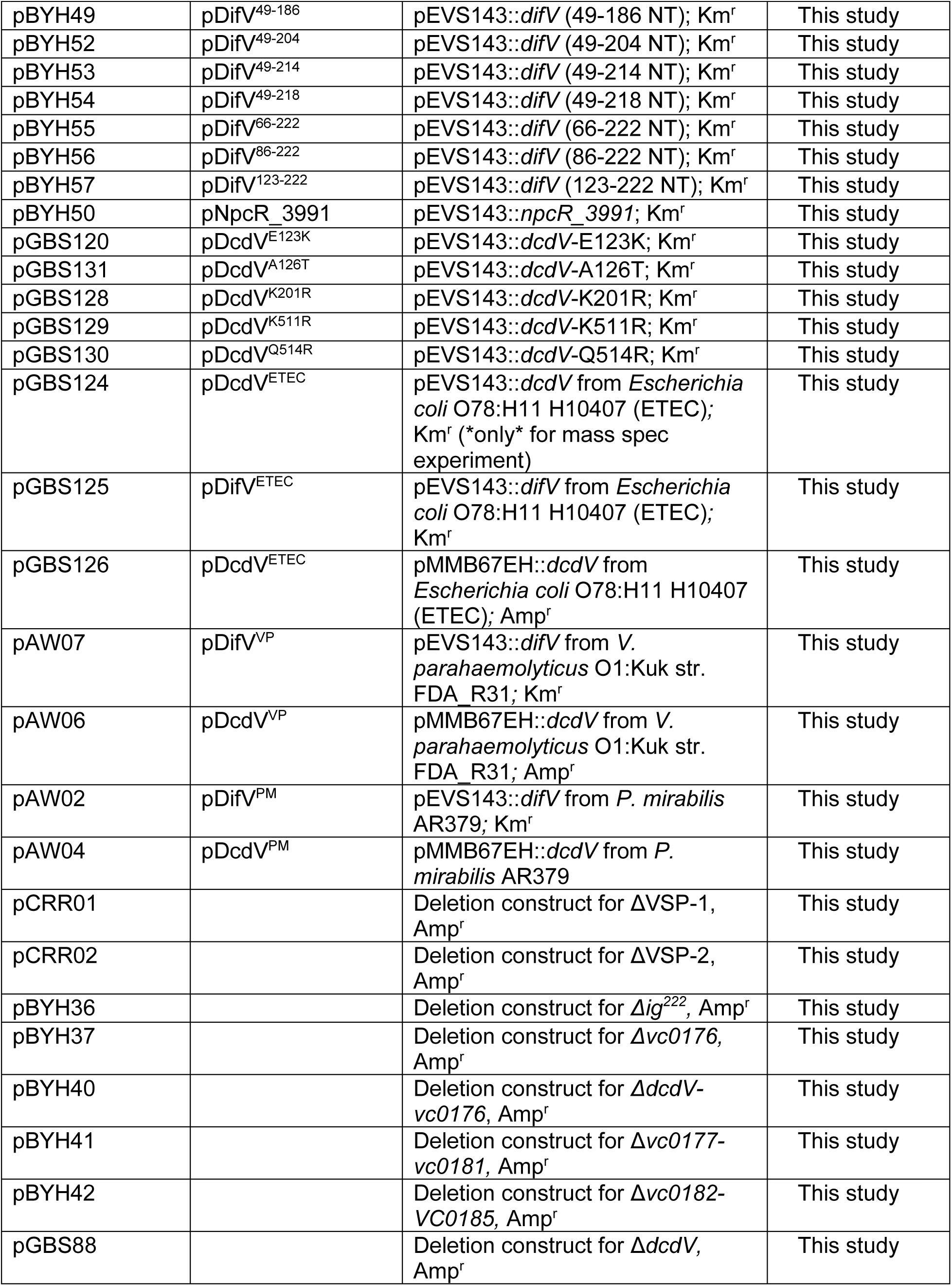
Plasmids Descriptions

**Supplementary Table 3.**
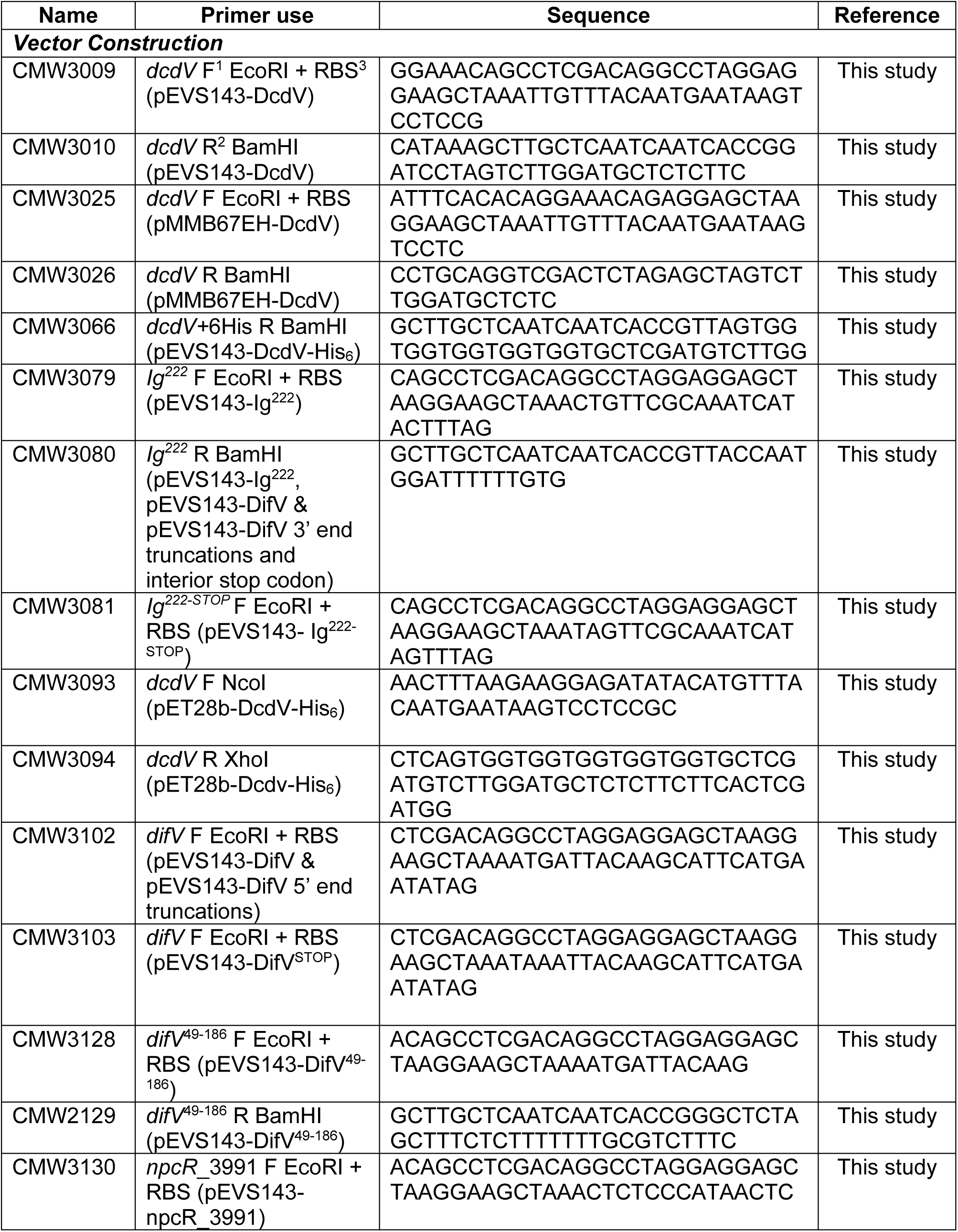
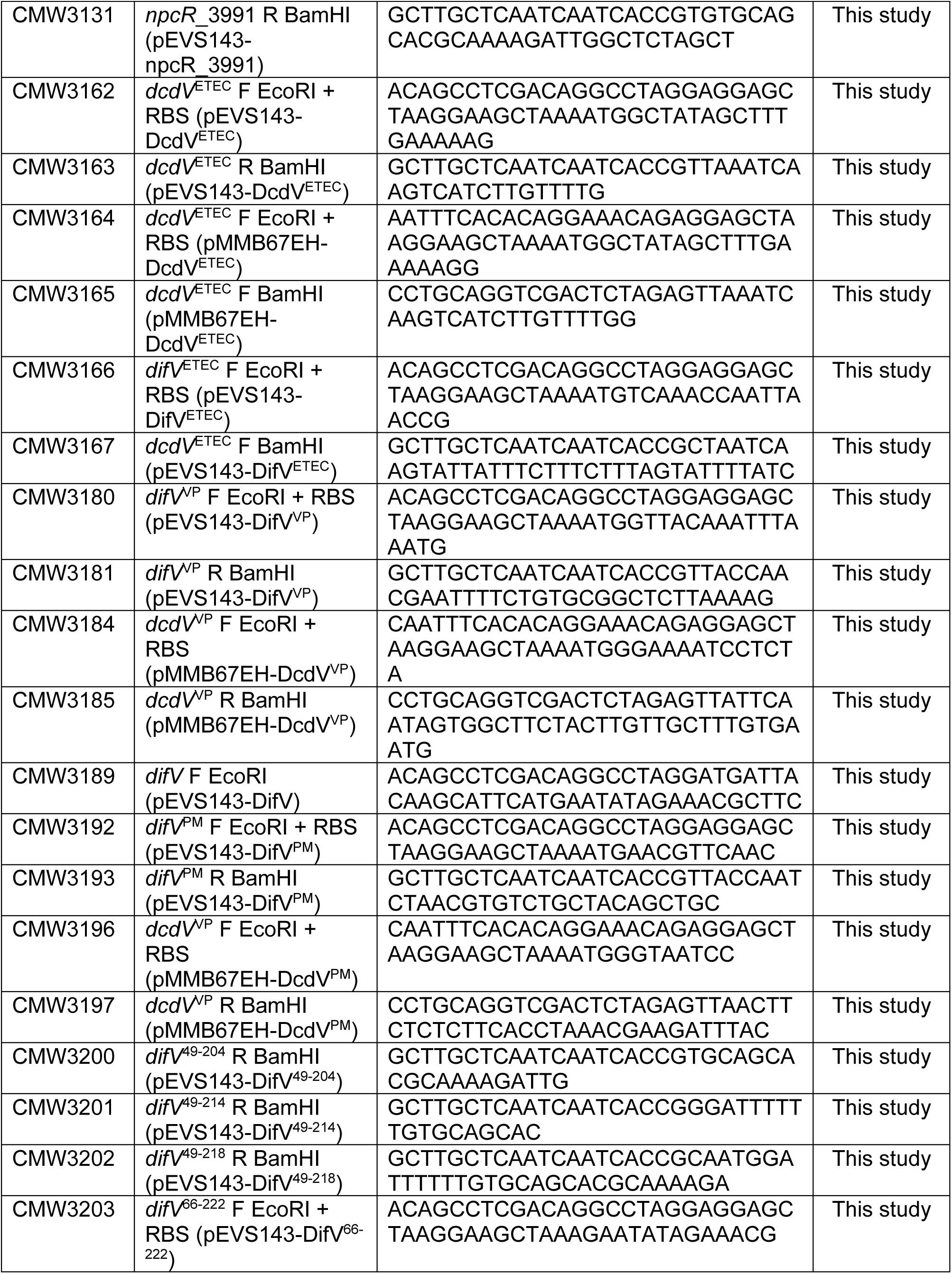
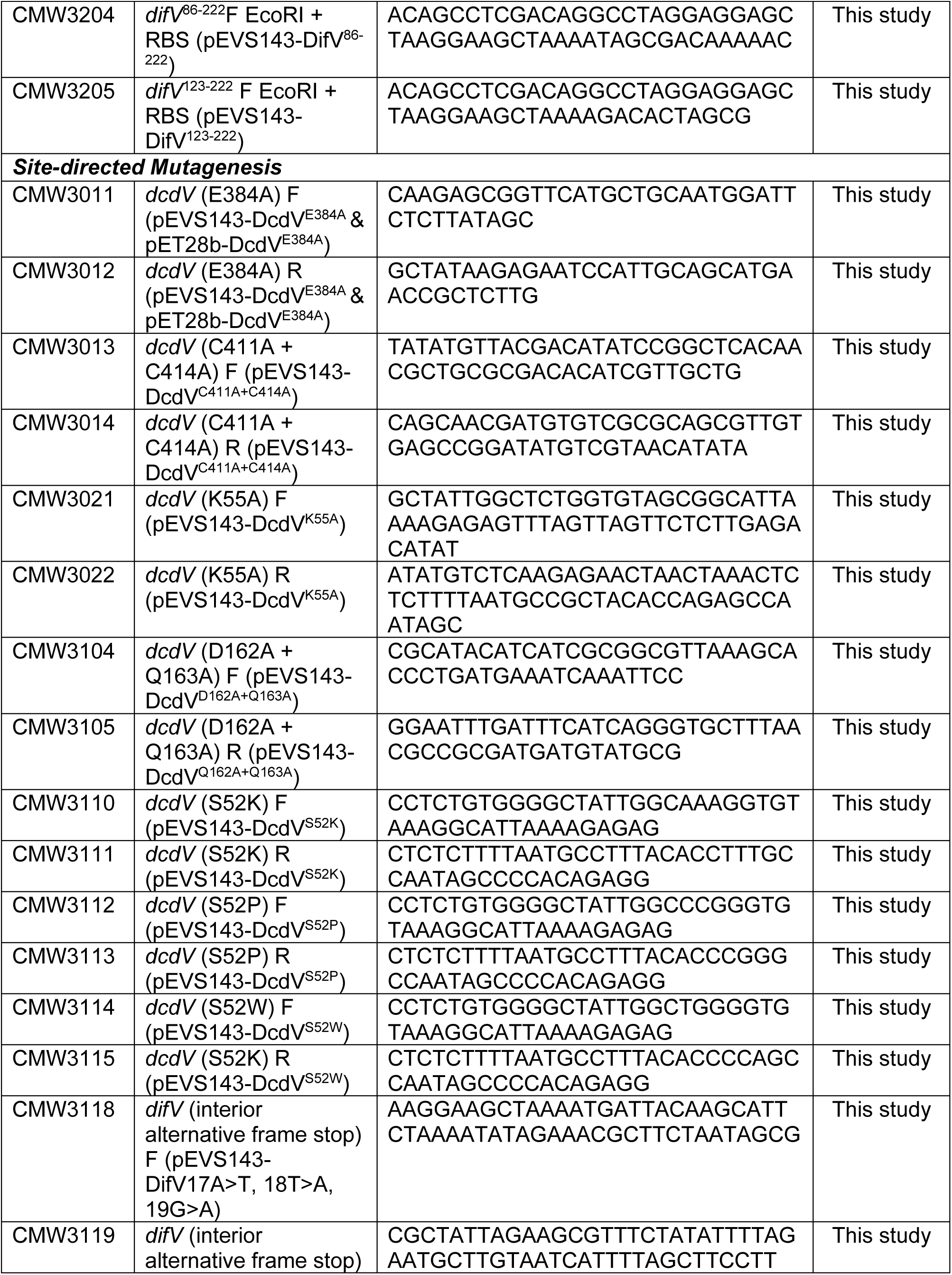
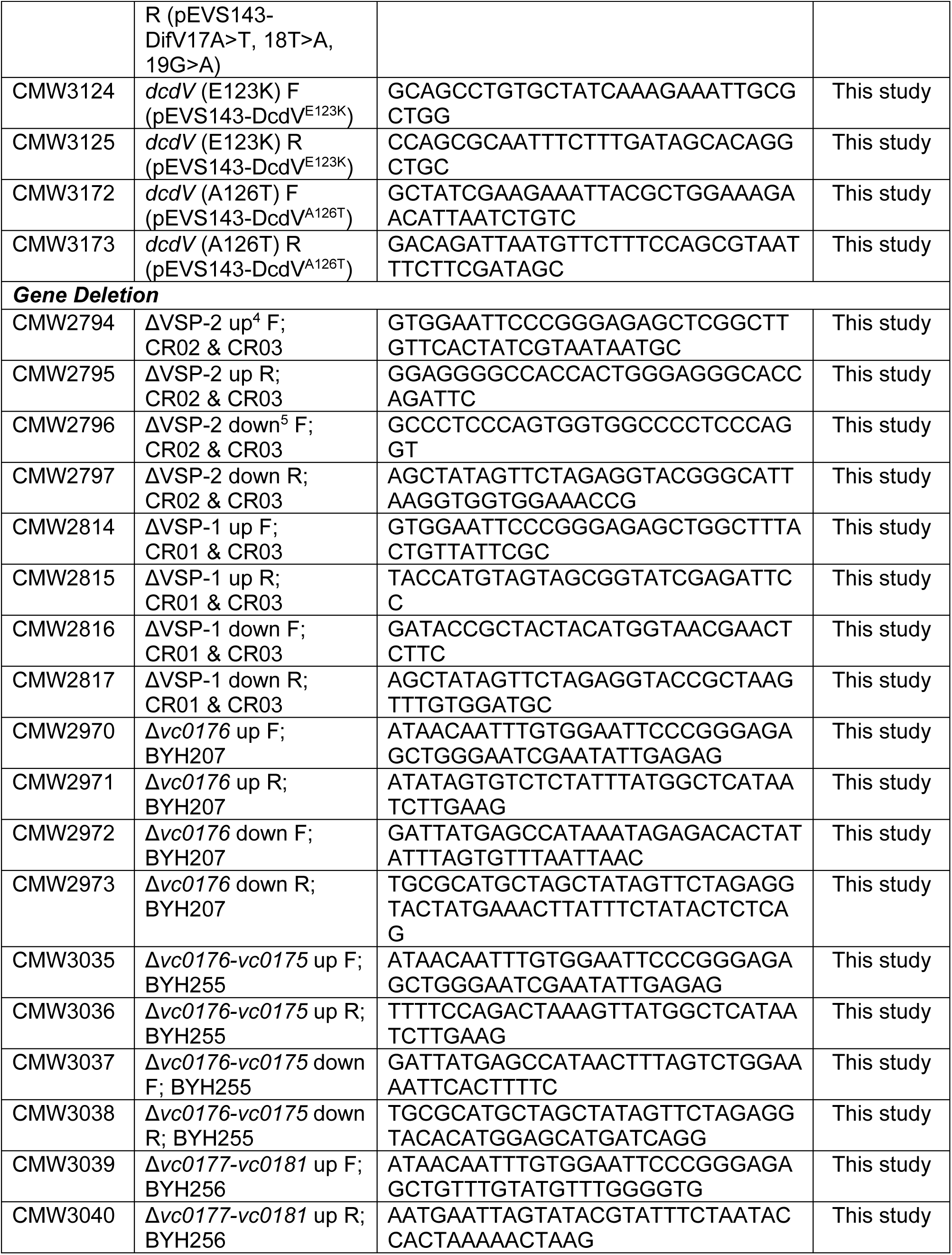
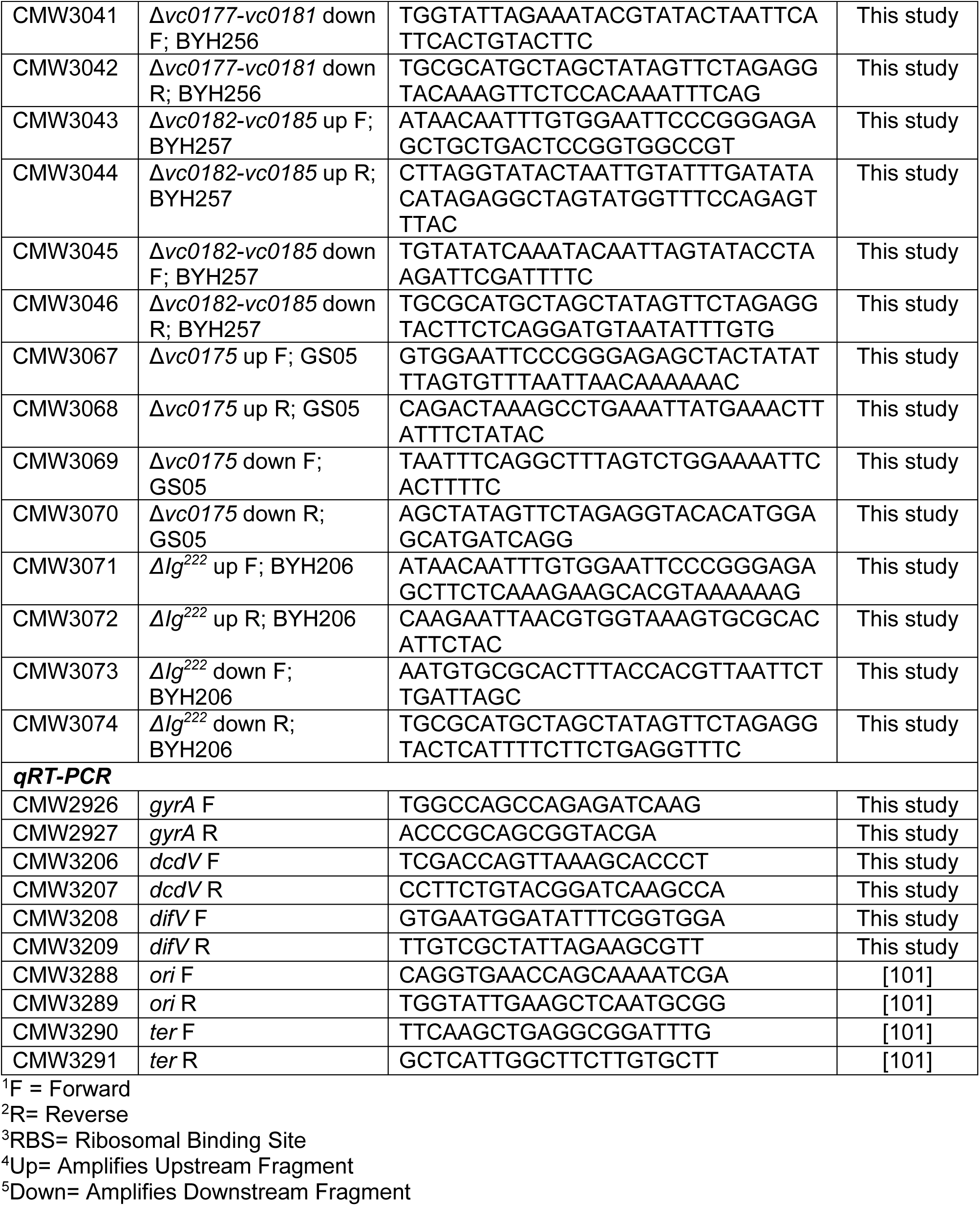
Oligonucleotides used in this study.

**Supplementary Table 4.** This table sorts the indicated lineages by the DcdV homolog in that group with the maximum amino acid similarity to *V. cholerae* DcdV.

| DcdV homologs summary table |  |  |
| --- | --- | --- |
| Lineages and percentage similarities of DcdV homologs containing both DCD and PLK domains |  |  |
| DomArch.Gene3D | Lineage | Max%Positive |
| PLK+DCD | Bacteria>Proteobacteria | 100.00 |
| PLK+DCD | Bacteria>Bacteroidetes | 58.80 |
| PLK+DCD | Bacteria>Balneolaeota | 56.02 |
| PLK+DCD | Bacteria>Actinobacteria | 55.63 |
| PLK+DCD | Archaea>Thaumarchaeota | 53.65 |
| PLK+DCD | Bacteria>Firmicutes | 52.27 |
| PLK+DCD | Bacteria>Planctomycetes | 52.27 |
| PLK+DCD | Bacteria | 51.88 |
| NABP+PLK+DCD | Bacteria>Proteobacteria | 51.09 |
| PLK+DCD | Bacteria>Acidobacteria | 49.70 |
| PLK+DCD | Bacteria>Verrucomicrobia | 48.69 |
| PLK+DCD | Bacteria>Chlamydiae | 45.25 |
| PLK+DCD+NABP | Bacteria>Proteobacteria | 42.48 |
| PLK+DCD+NABP+NABP | Bacteria>Proteobacteria | 39.85 |
| PLK+PLK+DCD | Bacteria>Proteobacteria | 36.75 |
| PLK+DCD | Bacteria>Cyanobacteria | 34.63 |
| PLK+DCD | Eukaryota>Ascomycota | 27.63 |
| PLK+DCD | Eukaryota>Ciliophora | 27.57 |
| PLK+DCD | Eukaryota>Basidiomycota | 25.10 |
| PLK+DCD | Eukaryota>Chytridiomycota | 23.77 |
| PLK+DCD | Eukaryota>Mucoromycota | 22.63 |
| PLK+DCD | Eukaryota>Apicomplexa | 19.96 |
| PLK+DCD | Eukaryota>Streptophyta | 19.39 |
| PLK+Znf_CCHC+DCD | Eukaryota>Ascomycota | 16.92 |

**Key** (% pos)
**Abbreviations.** PLK, P-loop containing nucleotide triphosphate hydrolases; DCD, Cytidine Deaminase domain 2; NABP, Nucleic acid-binding proteins; Znf\_CCHC, Zinc finger CCHC-type

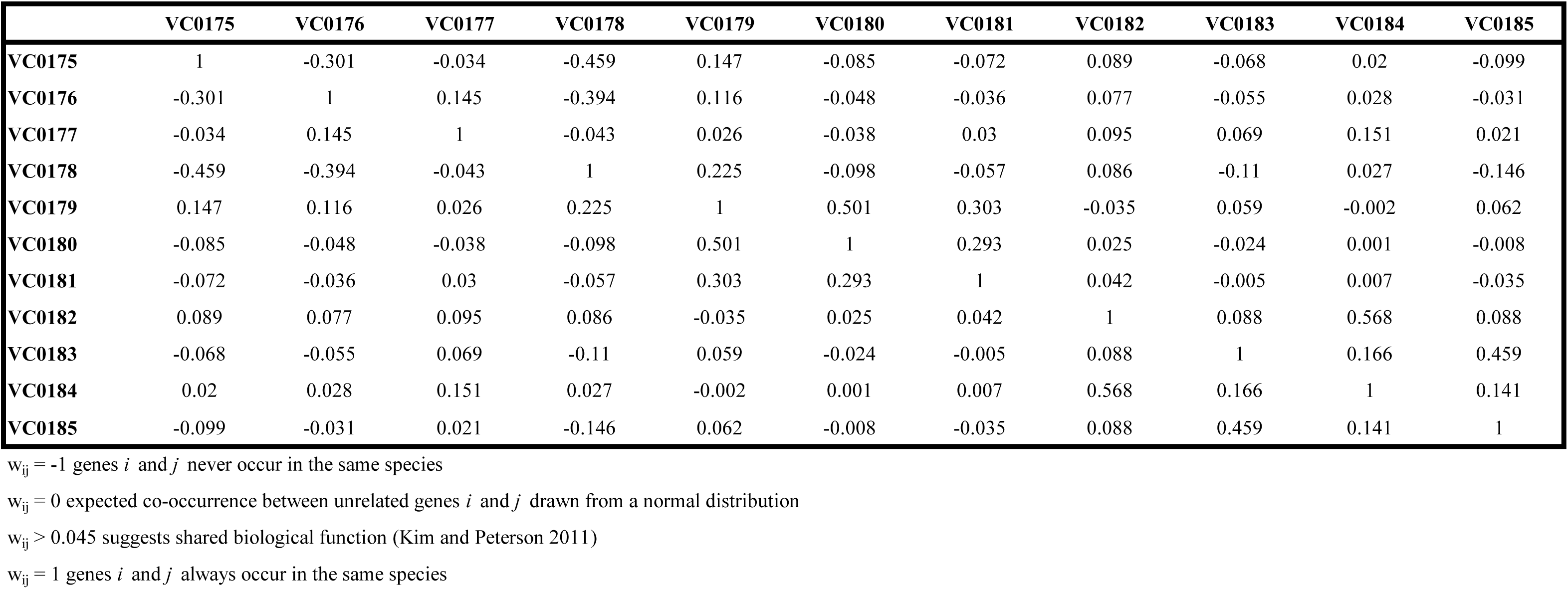
Partial Correlation Value w_ij_ of VSP-1 Genes *i* to *j* (Supplemental File 1)

|  | VC0175 | VC0176 | VC0177 | VC0178 | VC0179 | VC0180 | VC0181 | VC0182 | VC0183 | VC0184 | VC0185 |
| --- | --- | --- | --- | --- | --- | --- | --- | --- | --- | --- | --- |
| VC0175 | 1 | -0.301 | -0.034 | -0.459 | 0.147 | -0.085 | -0.072 | 0.089 | -0.068 | 0.02 | -0.099 |
| VC0176 | -0.301 | 1 | 0.145 | -0.394 | 0.116 | -0.048 | -0.036 | 0.077 | -0.055 | 0.028 | -0.031 |
| VC0177 | -0.034 | 0.145 | 1 | -0.043 | 0.026 | -0.038 | 0.03 | 0.095 | 0.069 | 0.151 | 0.021 |
| VC0178 | -0.459 | -0.394 | -0.043 | 1 | 0.225 | -0.098 | -0.057 | 0.086 | -0.11 | 0.027 | -0.146 |
| VC0179 | 0.147 | 0.116 | 0.026 | 0.225 | 1 | 0.501 | 0.303 | -0.035 | 0.059 | -0.002 | 0.062 |
| VC0180 | -0.085 | -0.048 | -0.038 | -0.098 | 0.501 | 1 | 0.293 | 0.025 | -0.024 | 0.001 | -0.008 |
| VC0181 | -0.072 | -0.036 | 0.03 | -0.057 | 0.303 | 0.293 | 1 | 0.042 | -0.005 | 0.007 | -0.035 |
| VC0182 | 0.089 | 0.077 | 0.095 | 0.086 | -0.035 | 0.025 | 0.042 | 1 | 0.088 | 0.568 | 0.088 |
| VC0183 | -0.068 | -0.055 | 0.069 | -0.11 | 0.059 | -0.024 | -0.005 | 0.088 | 1 | 0.166 | 0.459 |
| VC0184 | 0.02 | 0.028 | 0.151 | 0.027 | -0.002 | 0.001 | 0.007 | 0.568 | 0.166 | 1 | 0.141 |
| VC0185 | -0.099 | -0.031 | 0.021 | -0.146 | 0.062 | -0.008 | -0.035 | 0.088 | 0.459 | 0.141 | 1 |

$w_{ij} = -1$ genes $i$ and $j$ never occur in the same species
$w_{ij} = 0$ expected co-occurrence between unrelated genes $i$ and $j$ drawn from a normal distribution
$w_{ij} > 0.045$ suggests shared biological function (Kim and Peterson 2011)
$w_{ij} = 1$ genes $i$ and $j$ always occur in the same species

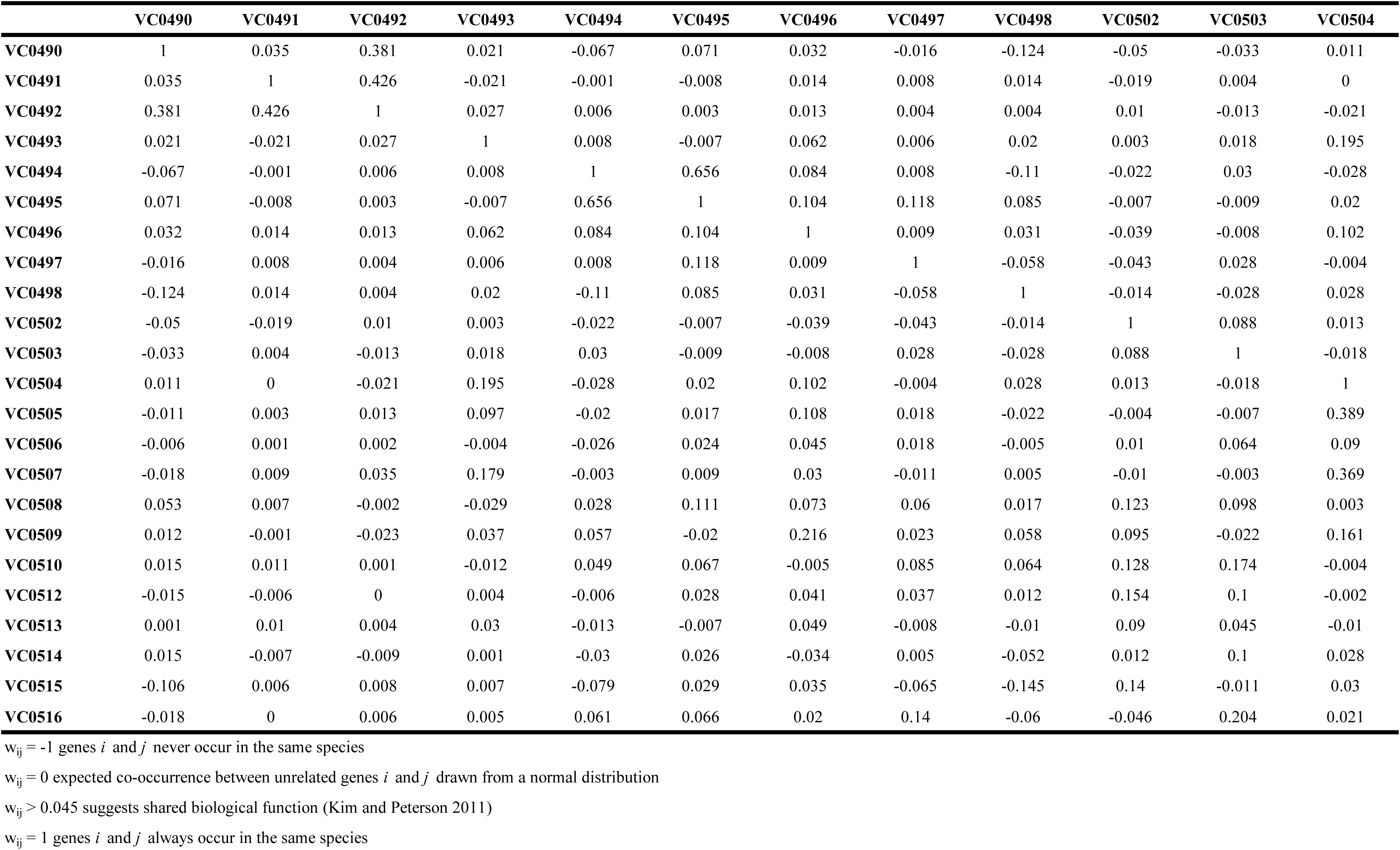
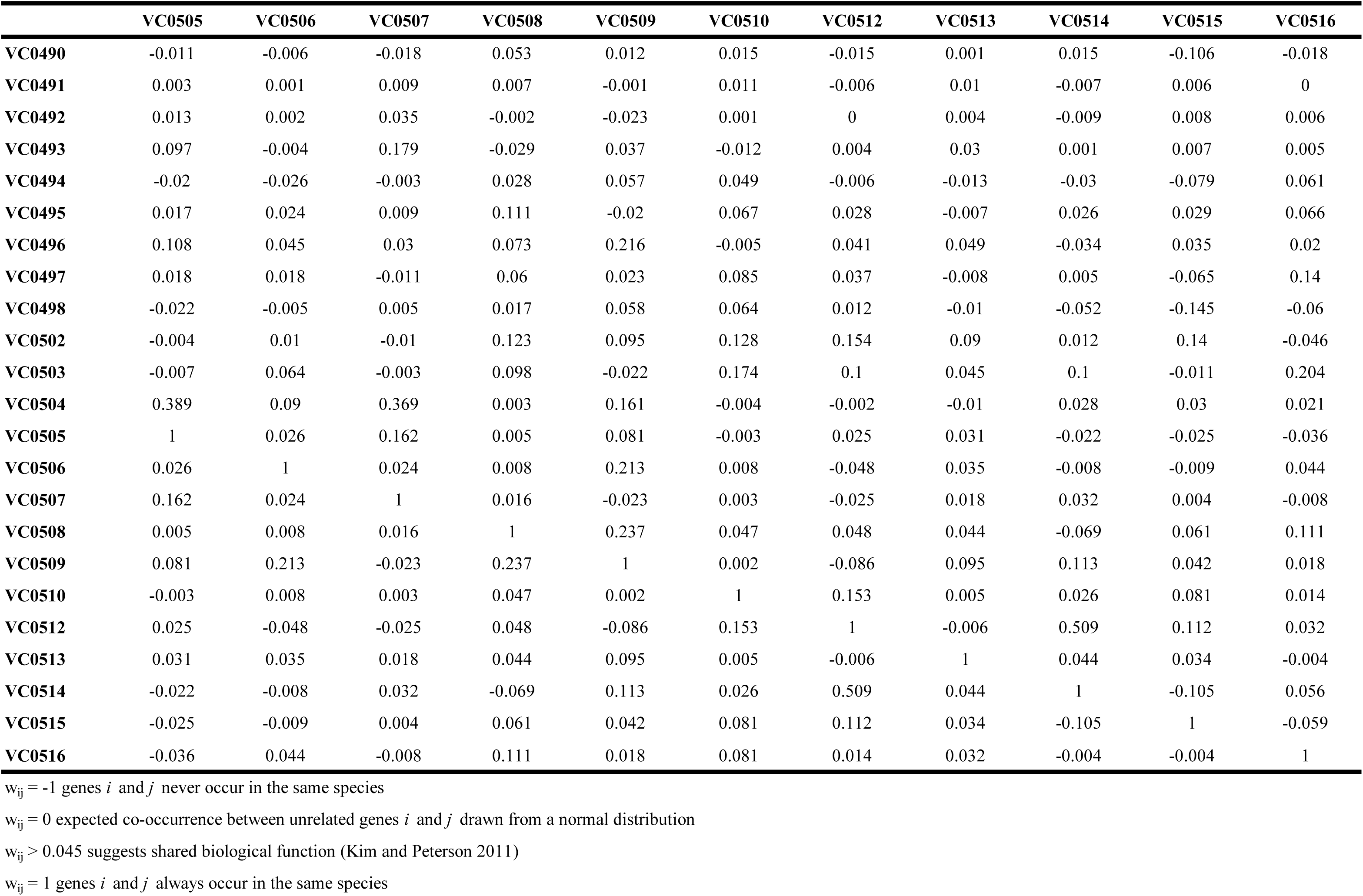
Partial Correlation Value w_ij_ of VSP-2 Genes *i* to *j* (Supplemental File 2)

